# Whole-genome scanning reveals selection mechanisms in epipelagic *Chaetoceros* diatom populations

**DOI:** 10.1101/2022.05.19.492674

**Authors:** Charlotte Nef, Mohammed-Amin Madoui, Éric Pelletier, Chris Bowler

## Abstract

Diatoms form a diverse and abundant group of photosynthetic protists that are essential players in marine ecosystems. However, the microevolutionary structure of their populations remains poorly understood, particularly in polar regions. Exploring how closely related diatoms adapt to different oceanic ecoregions is essential given their short generation times, which may allow rapid adaptations to different environments; and their prevalence in marine regions dramatically impacted by climate change, such as the Arctic and Southern Oceans. Here, we address genetic diversity patterns in *Chaetoceros*, the most abundant diatom genus and one of the most diverse, using 11 metagenome-assembled genomes (MAGs) reconstructed from *Tara* Oceans metagenomes. Genome-resolved metagenomics on these MAGs confirmed a prevalent distribution of *Chaetoceros* in the Arctic Ocean with lower dispersal in the Pacific and Southern Oceans as well as in the Mediterranean Sea. Single nucleotide variants identified within the different MAG populations allowed us to draw a first landscape of *Chaetoceros* genetic diversity and to reveal an elevated genetic structure in some Arctic Ocean populations with F_ST_ levels ranging up to ≥ 0.2. Genetic differentiation patterns of closely related *Chaetoceros* populations appear to be correlated with abiotic factors rather than with geographic distance. We found clear positive selection of genes involved in nutrient availability responses, in particular for iron (e.g., ISIP2a, flavodoxin), silicate and phosphate (e.g., polyamine synthase), that were further confirmed in *Chaetoceros* transcriptomes. Altogether, these results provide new insights and perspectives into diatom metapopulation genomics through the integration of metagenomic and environmental data.

## Introduction

About half of primary productivity on Earth is supported by aquatic phytoplankton, a phylogenetically diverse group of photosynthetic organisms composed of eukaryotic algae and cyanobacteria that provide essential ecosystem services, from nutrient cycling and CO2 regulation to sustaining superior trophic levels as the base of marine food webs (Nelson et al. 1995; Field et al. 1998; Falkowski et al. 2004). Among phytoplankton, diatoms are pivotal in marine ecosystems since they account for an estimated 40% marine primary productivity and 20% global carbon fixation (Field et al. 1998), as well as being important contributors to global carbon export (Guidi et al. 2016). Moreover, they link silicon and carbon biogeochemical cycles through the synthesis of their elaborate silicified cell wall, surrounded and embedded by glycoproteins that prevent its dissolution (Lewin 1961; Kröger and Sumper 1998). Diatoms are therefore key players also in the global silicon cycle, particularly in the Southern Ocean (Tréguer and de La Rocha 2013; Llopis Monferrer et al. 2021).

Like other pelagic plankton, diatoms are thought to display high dispersion potential due to their rapid generation times and large population sizes, combined with the few apparent oceanic barriers to dispersal (Norris 2000; Cermeño and Falkowski 2009). As a consequence, they are expected to show reduced diversity patterns and biogeographic structure due to homogenised genetic pools (Finlay 2002). Instead, molecular surveys have revealed that diatom populations exhibit tremendous diversity, with more than 4,000 different operational taxonomic units (OTUs) (de Vargas et al. 2015), while being widely distributed across all major oceanic provinces (Malviya et al. 2016) encompassing high latitudes, upwelling regions as well as stratified waters (Kemp and Villareal 2018; Leblanc et al. 2018). The ecological success of diatoms is undoubtedly linked to their complex evolutionary history, which was found to be sustained by horizontal gene transfers from bacteria (Armbrust et al. 2004; Bowler et al. 2008), and mosaic plastid evolution derived from both red and green algae (Moustafa et al. 2009; Dorrell and Smith 2011; Dorrell et al. 2017). This chimeric origin led to specific physiological innovations, such as silicon utilisation for cell protection, efficient nutrient uptake systems allowing rapid responses to environmental fluctuations, a functional urea cycle and potential carbon concentration mechanisms (Armbrust et al. 2004; Bowler et al. 2008; Pierella Karlusich, Bowler, et al. 2021). In contrast to these genetically-encoded functions, diatom genomes themselves appear to display a wide variety of dynamics, through specific transposable elements in the model diatoms *Thalassiosira pseudonana* and *Phaeodactylum tricornutum* (Armbrust et al. 2004; Maumus et al. 2009), alternative splicing (Rastogi et al. 2018), as well as gene copy number variation and mitotic recombination between homologous chromosomes (Bulankova et al. 2021). Altogether, these characteristics likely fuel diatom diversity, leading to rapid diversification rates (Bowler et al. 2008) while increasing their ability to respond to changing environmental conditions.

Climate change is expected to induce a range of environmental stressors on phytoplankton (Hays et al. 2005). Among these are increased water temperature and stratification, nutrient paucity and acidification (Bopp et al. 2013). Moreover, a recent study indicated that numerous important diatom genera, such as *Chaetoceros, Porosira* and *Proboscia*, are predicted to be vulnerable to climate change, particularly in polar plankton communities (Chaffron et al. 2021). Diatoms appear therefore to be valuable candidates to investigate the fundamental links between their genomes, physiology and population dynamics, in light of predicted environmental changes. Understanding such principles would require access to the genome of natural diatom populations as well as precise contextual information. With the emergence of new sequencing technologies and processes to recover genomes from environmental data, either from metagenomes or single-cell genomes, it is now possible to access the genomic information of organisms by going beyond culture-dependent approaches, allowing us to gain insights into the biology and ecology of natural populations (Iverson et al. 2012; Mangot et al. 2017). This is of particular interest for organisms for which culture conditions cannot be mimicked easily, as for instance organisms thriving in polar environments. These new techniques have enabled the scientific community to access sequences from taxa lacking significant information, such as Euryarchaeota (Iverson et al. 2012), Picozoa (Not et al. 2007; Seenivasan et al. 2013), MOCHs (for Marine OCHrophytes) (Massana et al. 2014), MAST-4 (for MArine STramenopiles) (Mangot et al. 2017), and rappemonads (Kim et al. 2011).

Among diatoms, the genus *Chaetoceros* holds a particular position as it is the most widespread diatom genus, presenting a worldwide distribution from pole to pole with a prevalence at high latitudes (Malviya et al. 2016; De Luca, Kooistra, et al. 2019; Sommeria-Klein et al. 2021). As such, it is considered an important driver of carbon export and silica sinking in modern oceans (Smetacek 1999; Tréguer et al. 2018). The genus displays a high level of diversity, with 239 accepted species names in Algaebase, in addition to 153 names under debate or yet to be verified (https://www.algaebase.org, as of May 2022). It is generally accepted that the *Chaetoceros* genus is subdivided into the *Hyalochaete* and *Phaeoceros* subgenera, the latter including the type species *Chaetoceros dichaeta*, though their exact subdivision remains under debate (De Luca, Sarno, et al. 2019). *Chaetoceros* presents peculiar physiological properties that may be responsible for its prevalent distribution. For instance, some *Chaetoceros* species have been shown to display unusually high C:N ratios unaffected by light regime and nitrogen source, suggesting a capacity to accumulate superior carbon per nitrogen units than other Arctic diatoms, while showing physiological responses similar to those of more temperate diatoms (Schiffrine et al. 2020). Besides its particular physiological characteristics, *Chaetoceros* is known to participate in a significant range of associations with a wide variety of microorganisms. The *Chaetoceros* phycosphere has been shown to gather a diverse set of epibiotic bacteria, the composition of which simplifies along subculturing (Crenn et al. 2018), and is significantly influenced by nutrient availability and host growth stage (Baker et al. 2016). Some associated bacteria have even been observed to favour resistance of *Chaetoceros* cells against viral infection and lysis compared to axenic controls (Kimura and Tomaru 2014). *Chaetoceros* can be involved in photosymbioses with epibiotic peritrich and tintinnid ciliates (Gómez 2020), interact with nitrogen-fixing cyanobacteria in diatom-diazotroph associations (Foster et al. 2011; Pierella Karlusich, Pelletier, et al. 2021) and is globally highly connected with other plankton members in the *Tara* Oceans network of planktonic associations (Vincent and Bowler 2020). Therefore, given the ecological significance of *Chaetoceros* and its prevalence in regions particularly predicted to be vulnerable to climate change, the present study focuses on describing patterns of genetic diversity and population structure of this diatom genus. To this end, we leveraged 11 metagenome-assembled genomes (MAGs) originating from the *Tara* Oceans expeditions (Delmont et al. 2022), and that are associated with highly contextualised metadata. We aimed to answer the following questions: How are natural *Chaetoceros* populations structured? Is geographic distance a barrier to gene flow and, if not, what main ecological factors are correlated with *Chaetoceros* micro-diversification? What are the genetic functions undergoing selection among different *Chaetoceros* populations?

## Results

### Description and comparative analysis of *Chaetoceros* MAGs

The MAGs (see Supplementary Table S1 for details on their names) displayed genome sizes ranging from 10.6 (ARC_232) to 44.4 (PSW_256) Mbp that are the same order of magnitude as the genomes of the model diatoms *T. pseudonana* and *P. tricornutum* (Fig. 1A). The MAG gene numbers, ranging from 5,000 to 17,000 genes, adequately mirrored the genome sizes (Fig. 1C). Overall, the genomes displayed good completion percentages with 7 out of the 11 MAGs having a BUSCO score at least equal to 50% (Fig. 1B). The average percentage of GC ranged from 39% (ARC_267) to 44% (SOC_37), which is lower than those of *T. pseudonana* and *P. tricornutum* (Fig. 1E), with a global decreasing percentage of GC from first to third position in the codons (Fig. 1F). Mean gene lengths varied between 400 and 500 bp (Fig. 1D), again lower than those of *T. pseudonana* and *P. tricornutum*, but expected because metagenome-assembled genomes are generally more fragmented than genomes sequenced from cultured organisms (Bowers et al., 2017). A principal component analysis (PCA) on eight genome and gene metrics was conducted to test whether the MAGs belonging to the same geographical region displayed common genome characteristics (Fig. 1G-H). The PCA showed three defined groups: one consisting of ARC_116, ARC_217, PSE_171, PSE_253 and SOC_37 that are characterised by a larger number of genes, large genome and gene sizes as well as higher GC content, indicating a more compact genome. A second group consisted of ARC_267 and PSW_256 that display the largest intron sizes, and a last one grouped ARC_232 and SOC_60, the smallest *Chaetoceros* MAGs. This analysis did not reveal any clustering of the MAGs based on their geographical origins. It must be noted that some of the differences observed regarding for instance genome size may be linked to the reconstruction methods applied rather than to *bona fide* biological differences, as exemplified by different genome completion levels (see Fig. 1B).

**Figure 1.**
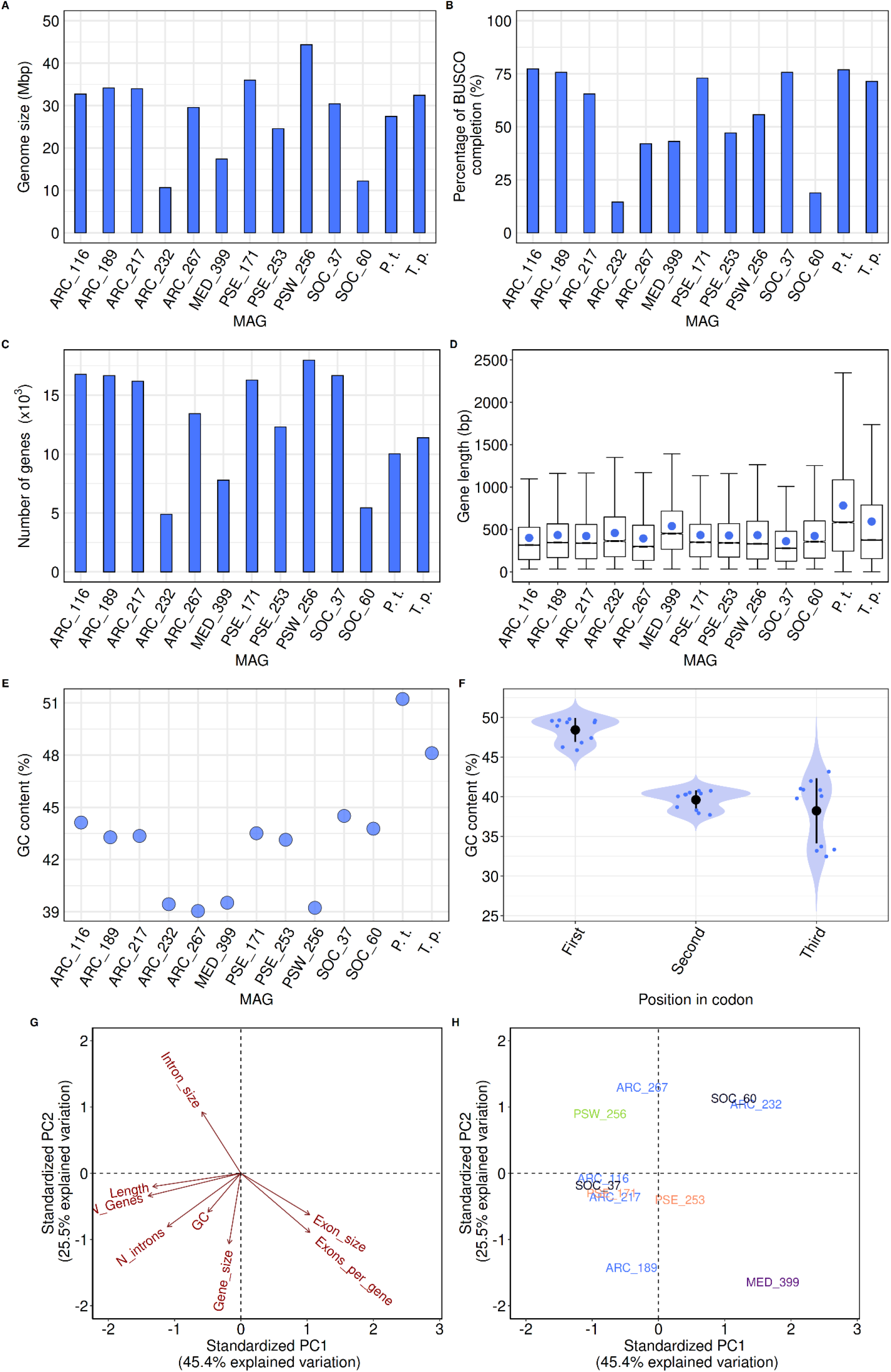
Characteristics of the different *Chaetoceros* MAGs. (A) Genome size, (B) level of BUSCO completion, (C) number of genes, and (D) boxplots of mean gene length (mean gene length is represented by the blue dot) of the MAGs and reference diatoms *P. tricornutum* (P.t.) and *T. pseudonana* (T.p.). (E) Mean GC content of MAGs and reference diatom genomes. (F) Distribution of GC content along codon positions of the MAGs. (G-H) Principal Component Analysis of different gene and genome metrics of the MAGs, shaded by geographical origin (blue: Arctic Ocean; purple: Mediterranean; orange: Pacific South Eastern Ocean; green: Pacific South Western Ocean; black: Southern Ocean).

Relatively weak average nucleotide identity (ANI) (< 80%) and average amino acid identity (AAI) (< 60%) were observed between the MAGs that were derived from the same geographical areas, suggesting that the populations are not necessarily closely related (Fig. 2A, Supplementary Figure S1A). Pairwise ANI and AAI values were highly positively correlated particularly for pairwise ANI > 85% and AAI > 75% (Supplementary Figure S1A-B). The most elevated ANI (95.1%) and AAI (94%) were observed between the MAGs ARC_217 and PSE_171. Elevated pairwise ANI (> 80%) and AAI (> 60%) values were also observed for PSE_253 and ARC_189 as well as for PSW_256 and ARC_267.

**Figure 2.**
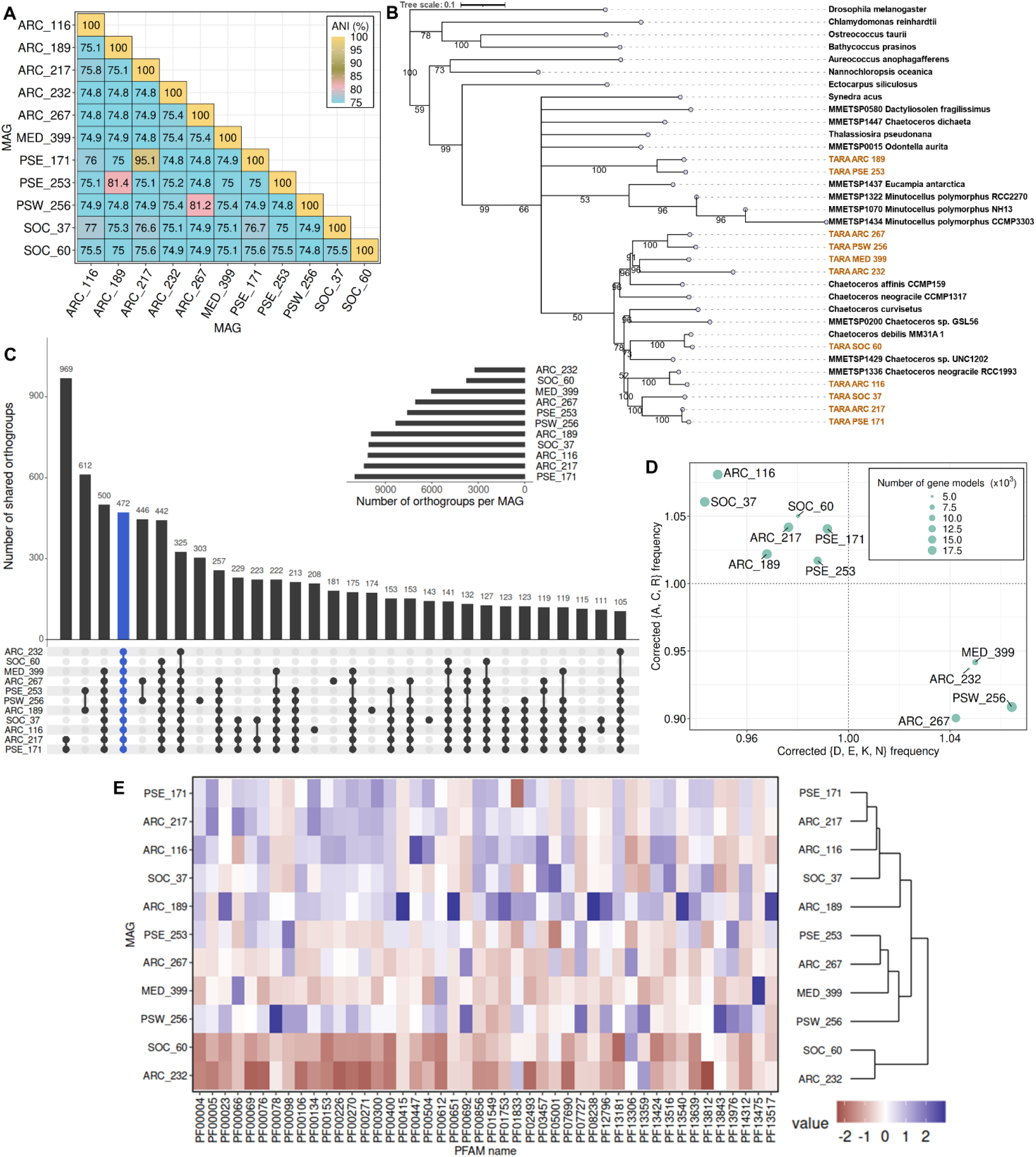
Comparative analysis of *Chaetoceros* MAGs and their coding potential. (A) Average Nucleotide Identity. (B) Concatenated multigene ML tree generated with RAxML (100 bootstrap), based on 83 BUSCO gene clusters with the *Chaetoceros* MAGs highlighted in gold. Bootstrap values ≥ 50% are indicated. (C) Upset plot representing the top 30 shared orthogroups among the MAGs, with the orthogroups shared by all genomes highlighted in blue. (D) Frequency of the most variable amino acids compared to their global means across all MAGs. The MAG respective number of genes is indicated for comparison. (E) Heatmap of 46 PFAM domains displaying the most variable copy number (SD ≥ 10) among the MAGs (see Supplementary Table S2 for details).

We then evaluated the relatedness of the *Chaetoceros* MAGs between one another and with respect to other taxa based on a concatenated tree of 34 taxa for 83 single-copy nuclear genes (total 42,525 amino acids) across the eukaryotic tree of life (Fig. 2B). We obtained a relatively good phylogeny of the taxa, with a monophyly of the diatoms. As observed previously, MAGs from a close geography did not appear to resolve together. In accordance with the ANI/AAI patterns, three MAG pairs resolved together with high support values, namely ARC_189 and PSE 253, ARC_217 and PSE_171, and ARC_267 and PSW_256 (Fig. 2B). Both ARC_189 and PSE_253 MAGs resolved at the level of *C. dichaeta*, which suggested that they belonged to the *Phaeoceros* subgenera. Conversely, the 9 other MAGs resolved in the same clade as *C. affinis, C. curvisetus* and *C. debilis*, suggesting closeness to the *Hyalochaete* subgenera. The MAGs ARC_267, PSW_256, ARC_232 and MED_399 resolved in clades close to *C. affinis* CCMP159. The MAG SOC_60 appears to be most closely related to *C. debilis* with high support (bootstrap value > 90%), while ARC_116 was closely related to the *C. neogracile* RCC1993 strain but not to *C. neogracile* CCMP1317.

Interestingly, identifying the orthologous genes shared between the MAGs showed that the most elevated number of orthogroups were not shared by the 11 genomes, as one would have expected, but rather by the two MAGs ARC_217 and PSE_171 (969 orthogroups) (Fig. 2C). The second highest set of orthogroups was shared between ARC_189 and PSE_253 (612 orthogroups). The third group consisted of all the genomes except ARC_232 and SOC_60 (500 orthogroups), suggesting that the two excluded MAGs are the most divergent. This pattern is consistent with the result of the multigene phylogeny, and may be partly explained by the fact that both these genomes are the smallest in size, number of genes and BUSCO completion (Fig. 1A-C). Another explanation would be an artifactual result due to low genome completion. A total of 472 orthogroups appeared common to the 11 genomes. MAG-specific orthogroup sets were retrieved, with PSW_256 displaying the highest number of MAG-specific orthogroups (303), which may be because this genome is the largest both in size and gene number (Fig. 1).

Clear discrepancies were observed regarding the proportions of amino acids encoded in the MAGs. A first group, composed of ARC_116-217 and all genomes associated to PSE and SOC, exhibited an enrichment in arginine (R) as charged residues, in alanine (A) for the hydrophobic ones as well as in cysteine (C); the other group, consisting of ARC_232 and 267, MED_399 and PSW_256, formed a monophyletic group, and showed a larger proportion of aspartate (D), glutamate (E) and lysine (K) as charged residues, and in asparagine (N) (Fig. 2D), suggesting a replacement of some residues displaying the same chemical properties. Such differences in amino acid composition of predicted proteomes have been previously identified in a study investigating more than 100 algal genomes, in which saltwater algae encoded higher proportions of D, E and K residues and lower proportions of A and C compared to freshwater species (Nelson et al. 2021).

We further conducted a comparative analysis at the level of PFAM domains to test the relative genome enrichment in putative biological functions. Most MAGs showed a median of two PFAM domains per gene, with ARC_116, ARC_189 and SOC_37 among the most complete MAGs, displaying 3 domains per gene (Supplementary Figure S2). A selection of the most variable PFAM domains among the MAGs was conducted on those displaying a standard deviation at least equal or superior to 10, leading to the identification of 46 PFAMs (see Supplementary Table S2 for detailed information) whose relative enrichments are represented on a heatmap (Fig. 2E). Again, no clear distinction between the MAGs based on their geographical localisation was observed. The group formed by ARC_232 and SOC_60 displayed globally the same patterns of PFAM enrichment and depletion compared to the other MAGs, with comparatively less domains, a pattern consistent with their smaller size. These MAGs were particularly depleted in chaperone associated domains (PF00004 and PF00226) and in an IQ calmodulin-binding motif (PF00612) involved in protein binding. ARC_232 showed a dramatic depletion of a pentatricopeptide repeats domain potentially involved in RNA metabolism (PF13812). A group gathered PSE_171, ARC_217, ARC_116, SOC_37 and ARC_189, most of them showing the same gene and genome characteristics. Compared to all the other MAGs, ARC_189 was dramatically enriched in domains associated with regulators of chromatin structure, domains containing repeat motifs and chromosome condensation repeats (PF00415, PF13540, PF08238, PF12796, PF13517 and PF00651). ARC_217 and PSE_171 where both enriched in the ATP-binding domain of ABC transporters and cyclins (PF00005 and PF00134) while the latter showed a strong depletion in PT/TIG domains (PF01833), a putative family of transcription factors. On the other hand, ARC_116 displayed a strong enrichment in heat shock transcription factor and chlorophyll a-b binding protein domains (PF00447 and PF00504) while SOC_37 displayed a significant enrichment in RNA polymerase Rpb1 C-terminal domain (PF05001). Another group consisted of the genomes PSE_253, ARC_267, MED_399 and PSW_256, all exhibiting medium BUSCO completion. Both PSE_253 and ARC_267 showed an enrichment in zinc-finger domain PF00098 while MED_399 was enriched in domains involved in ubiquitination processes (PF00066 and PF13475). Finally, of all the MAGs, PSW_256 was found strongly enriched in reverse transcriptase (PF00078) and transposase IS4 (PF13843, PiggyBac transposon) domains, a pattern that appeared consistent since this genome was the largest.

### Genome-resolved biogeography of *Chaetoceros*

The biogeographical distribution of the MAGs was investigated by estimating the proportion of *Tara* Oceans metagenomic reads that mapped on the eleven genomes. After filtration of the reads based on their identity and coverage (see Materials and Methods, Supplementary Figures S3 and S4 for details), a final number of 20 different sampling stations and/or depths were conserved. The *Chaetoceros* MAGs together recruited 0.71% of reads from these stations (all size fractions combined) and presented an amphitropical distribution, with a large prevalence in the Arctic Ocean and minor dispersal in the Pacific and Southern Oceans, as well as in the Mediterranean Sea. The relative contribution of the MAGs to the total metagenomic reads ranged from 0.03% for SOC_60 in the Southern Ocean to a local maximum of up to 4% for ARC_116 in the Arctic Ocean (Supplementary Table S3A-B). For a given depth, some stations (i.e., TARA_92, TARA_173, TARA_188, TARA_189, TARA_194, TARA_201 and TARA_205) appeared to harbour two and up to three different *Chaetoceros* MAGs (Fig. 3A), which suggests that our approach was precise enough to discriminate a mixture of populations from strains that are expected to be closely related. The four MAGs ARC_116, ARC_217, ARC_232 and SOC_37 were retrieved at both the surface and the deep-chlorophyll maximum (DCM) of the water column, with ARC_116 being the most widespread and dominant *Chaetoceros* MAG at the surface and SOC_37 at the DCM. The six MAGs ARC_189, ARC_267, PSE_171, PSE_253, PSW_256 and SOC_60 were found only at the surface of the water column while MED_399 was the only one retrieved solely at the DCM. Of note, SOC_37 was expected to be associated with both the Arctic and Southern Oceans (see Supplementary Table S4 in Delmont et al. (2022)), but it appeared to be restricted only to the Arctic Ocean in our analysis, likely due to the stringency of our filtration parameters. The co-occurrence patterns of the MAGs were addressed by performing pairwise correlation tests between the eleven MAGs using their metagenomic abundance. It appeared that none of the MAGs displayed significant co-occurrences or mutual exclusions, except for MAGs PSE_171 and PSE_253 (*p-value* = 0) that were associated with the same unique station (TARA_92) (Fig. 3B). Each of the different MAGs appeared restricted to a distinct environment characterised by a narrow range of temperature between 1 and 4 °C for the same MAG (Fig. 3C). Conversely, most of the MAG populations appeared to be distributed across a rather large spectrum of iron, silicate, phosphate and nitrate concentrations (Supplementary Figure S5).

**Figure 3.**
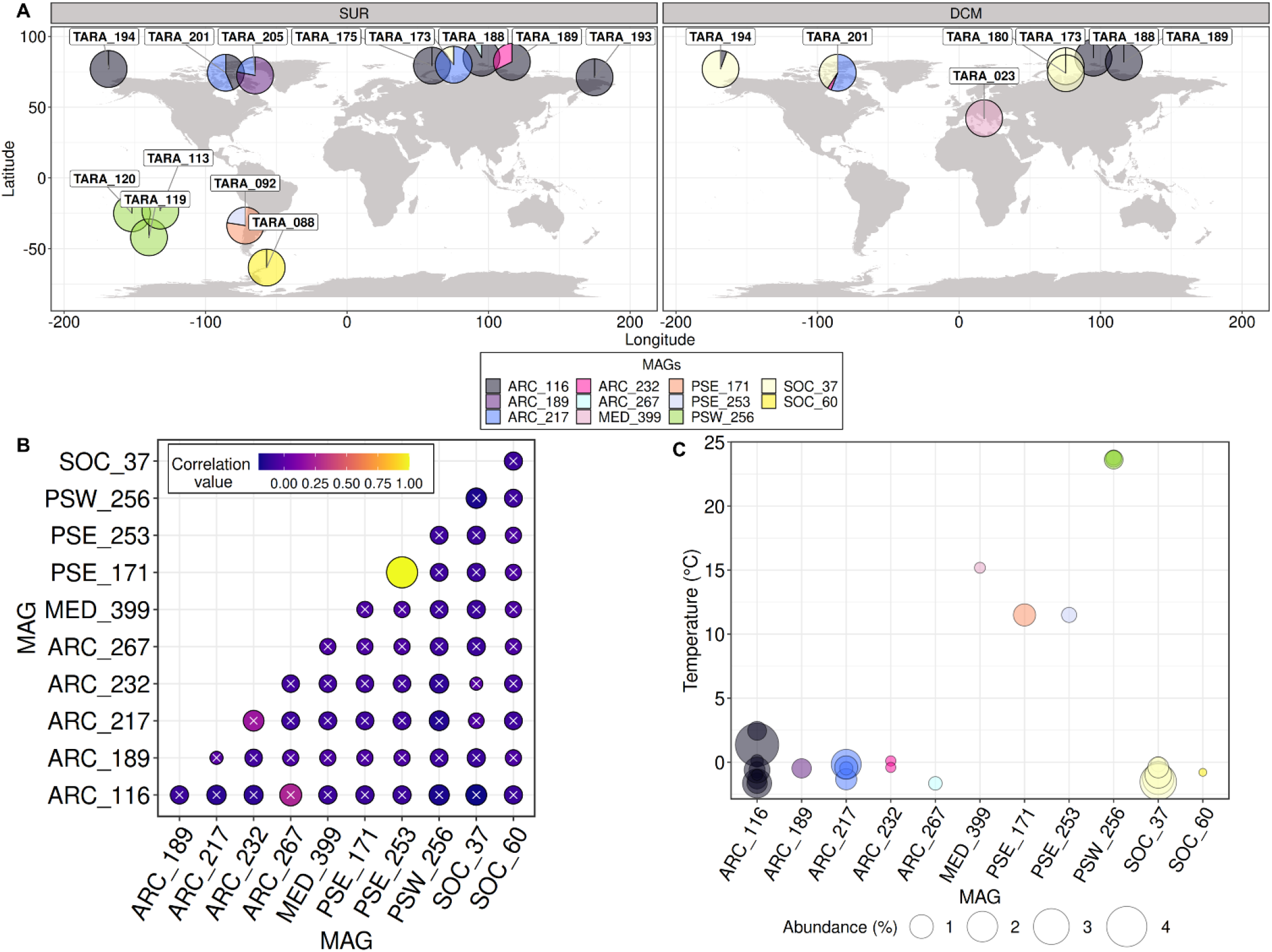
Biogeography of the *Chaetoceros* MAGs throughout *Tara* Oceans sampling sites. (A) Relative contributions of the *Chaetoceros* MAGs in surface (SUR) and deep-chlorophyll maximum (DCM) depths. (B) Pairwise correlation patterns of the MAGs (Pearson’s correlation *rho* shaded if superior to 0.05). (C) Bubble plot corresponding to the measured temperature at the sampling stations where each MAG was detected.

### Investigating genomic differentiation between *Chaetoceros* MAGs

#### *Chaetoceros* SNV landscape

We then investigated the level of genomic diversity in the different *Chaetoceros* populations by identifying for each MAG their respective single nucleotide variants. The number of variants of the *Chaetoceros* populations ranged from 1e5 to ~ 8e5 (Fig. 4B, Supplementary Figure S6), accounting for a total of 8,425,600 variants recruited. Globally, no significant correlation between the genome coverage and number of variants retrieved was observed (Pearson’s correlation *rho* = −0.22; *p*-value = 0.26) (Fig. 4A, Supplementary Table S4). Some MAGs, such as ARC_217, ARC_232 and PSW_256 nonetheless displayed variant patterns that followed the number of reads. Consequently, we assumed that the number of variants did not necessarily follow genome coverage and was rather dependent on the genome considered. The highest local SNV level ranged from 0.63% for ARC_217 at station TARA_205 (SUR) to as much as 2.34%, observed for ARC_116, which exhibited the highest range in terms of SNV levels, at station TARA_194 (DCM) in the Arctic Ocean (Fig. 4B, Supplementary Figure S6). This suggests that the average nucleotide identity of each MAG population to its respective consensus genome ranged between about 98% and 99%, a strong indicator illustrating the occurrence of local species harbouring non-negligible micro-diversity traits in different populations. We did not observe a significant correlation between the amount of SNVs and latitude but we noticed a rather strong correlation between SNVs and longitude (Supplementary Figure S7), which might be explained by the effect of oceanic currents in the Arctic Ocean. The most elevated mean SNV level depending on the MAG oceanic regions was observed for the *Chaetoceros* populations in the Pacific South Eastern Ocean (1.21%), followed by those in the Arctic Ocean (1.16%), the Southern Ocean (0.81%) and the Mediterranean (0.76%). Transition to transversion ratios ranged from 1.31 (ARC_217) to 1.96 (PSW_256), with a global average of 1.50 (Supplementary Figure S8). Overall, most of the *Chaetoceros* population variants were observed in the coding regions (49.28% mean value), followed by the intergenic (32.54%), UTR (14.11%) and intronic (4.07%) regions, a pattern consistently observed independently of the genome considered (Supplementary Figure S9). The variant effects were mostly missense mutations (53.29% mean value), followed by silent ones (45.97%) and a slight proportion of nonsense mutations (0.74%) (Supplementary Figure S10).

**Figure 4.**
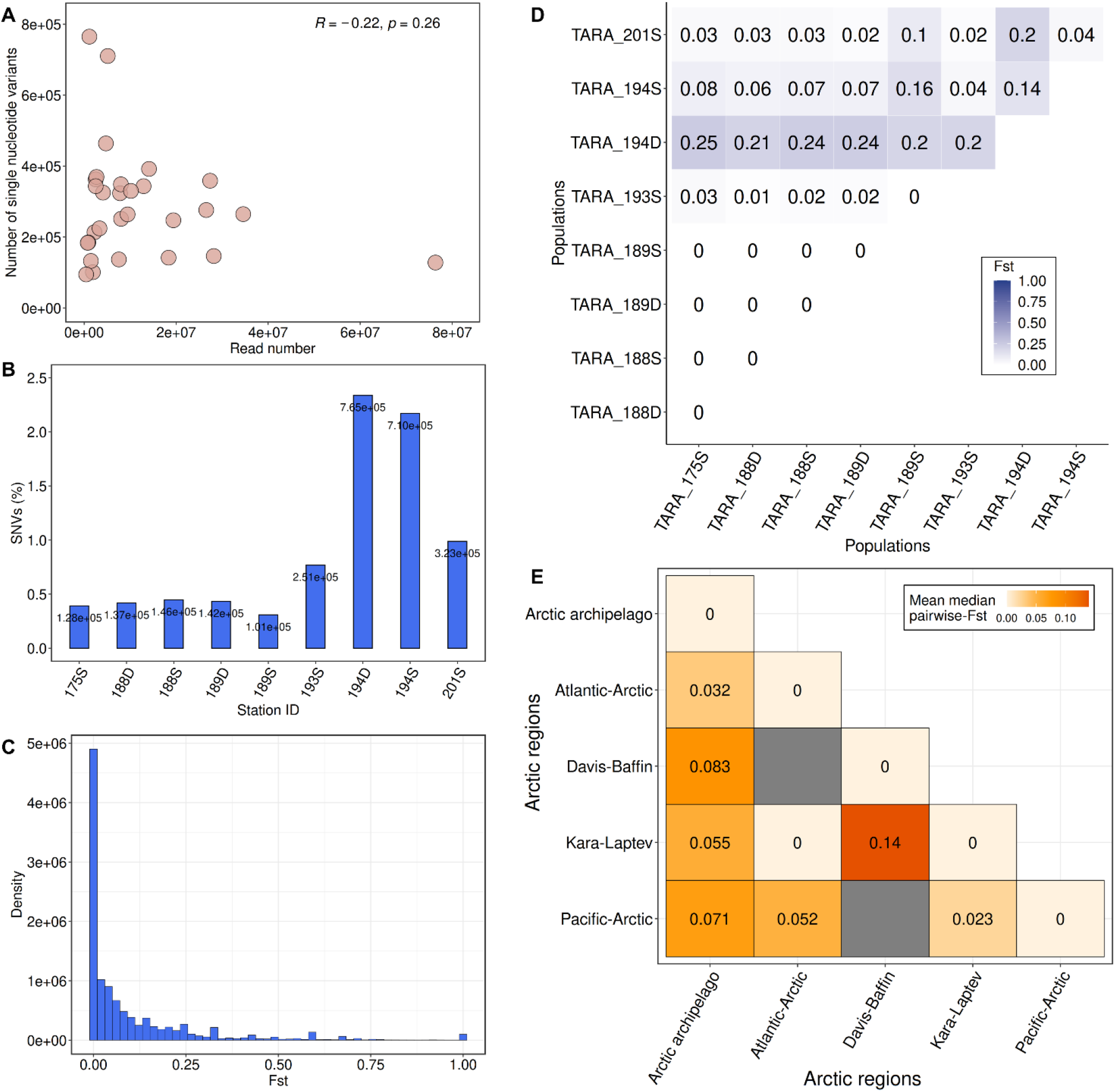
Population genomic analyses of *Chaetoceros* MAGs. (A) Scatterplot representing the number of SNVs compared to the number of reads for all samples considered in this study with Pearson’s correlation *rho* (n = 29). (B) Relative number of SNVs within ARC_116 populations. (C) *F*_ST_ distribution profile of ARC_116. (D) Pairwise *F*_ST_ matrix of ARC_116 populations. (E) Global pairwise *F*_ST_ matrix of all MAGs among Arctic Ocean regions (refers to ARC_116, ARC_189, ARC_217, PSW_256 and SOC_37). D: deep-chlorophyll maximum; S: surface.

#### Analysis of *Chaetoceros* population structure

We then investigated the level of population structure of the MAGs using the previously identified SNVs associated with the different populations and computed their pairwise fixation index (F-statistic or *F*ST). This index, which can range from 0 (no genetic differentiation) to 1 (complete differentiation), measures the extent of genetic inbreeding between populations using allele frequency, and is thus a proxy of their genetic distance (Wright 1965; Wright 1984). Among the detected variable loci, we selected the SNVs associated with the different populations for the MAGs that were present in at least two different sampling points (*Tara* Stations and/or depths), i.e., for ARC_116 (9 samples), ARC_217 (4), ARC_232 (2), PSW_256 (3) and SOC_37 (5). Plotting the population-wide *F*_ST_ distributions revealed globally unimodal patterns, indicative of a single species for each MAG (Fig. 4C, Supplementary Figure S11). ARC_116, which was the largest extant MAG, showed populations from stations TARA_175, TARA_188 and TARA_189 appearing to be genetically similar (pairwise *F*_ST_ of 0), consistent with their respective distance, and indicating that they formed one homogenous population (Fig. 4D). Noticeably, this MAG showed great (≥ 0.15) genetic differentiation at the DCM of station TARA_194, according to Wright’s guidelines for analysing bi-allelic loci (Wright 1984), and surface population at the same station also displayed pairwise *F*ST values distinct from the others but of lower magnitude, ranging from moderate to high (~0.05-0.15) genetic differentiation. This difference with the other ARC_116 populations may be at least partly explained by local marine currents, as station TARA_194 is influenced by inflow waters from the Pacific Ocean through the Bering Strait, and TARA_193 is enriched in cold waters circulating back to the Pacific Ocean. As evidenced by a pairwise *F*_ST_ of 0.14, both TARA_194 depths appeared to have moderate genetic differentiation among one another. This result suggested that this *Tara* station displayed rather important stratification patterns, resulting in the genetic differentiation of two sub-populations. Examining the metadata associated with this station revealed that the DCM was sampled 30 m deeper (35 m) than the surface (5 m) (Supplementary Table S5). Moreover, this sampling point showed distinct patterns of oxygen concentration and salinity between the depths as well as a phosphate enrichment at the DCM. The MAG ARC_217 showed relatively low pairwise *F*ST values indicating elevated connectivity between the populations, except between stations TARA_205 and TARA_173 (both SUR) located in the Davis-Baffin Bay and in the Kara-Laptev Seas (Supplementary Figures S12 and S13A). Both genomes ARC_232 and PSW_256 showed genetically similar populations (Supplementary Figure S13B-C), with PSW_256 exhibiting populations in stations TARA_113, TARA_119 and TARA_120 equally distinct from one another genetically. This pattern might be linked to these *Tara* stations being located between the Gambier Island archipelago in French Polynesia. Finally, the SOC_37 genome displayed globally low genetic differentiation, albeit slightly more elevated when compared with station TARA_173 at the surface than with the others. No clear differentiation with station TARA_201 was observed, although it is located at the opposite side on the Davis-Baffin Bay (Supplementary Figures S12 and S13D), suggesting a low effect of dispersal on their connectivity.

#### Global population structure among Arctic Ocean regions

We further compared the genomic differentiation of *Chaetoceros* populations between the Arctic regions, which were divided into five groups depending on their localisation: Pacific-Arctic, Kara-Laptev, Atlantic-Arctic, Arctic archipelago and Davis-Baffin. The most elevated genomic differentiation was between the Kara-Laptev and Davis-Baffin regions, which consistently appear opposite from one another (Fig. 4E; Supplementary Figure S12). The *Chaetoceros* populations located in the Arctic archipelago displayed globally moderate genetic differentiation compared to those in the other regions, with the largest difference being with the Davis-Baffin population. Low genetic structure was observed compared to the populations from the Atlantic-Arctic and Pacific-Arctic, both of which are located in zones with water influx from either the Atlantic or Pacific Oceans. Low differentiation was also noted between the Kara-Laptev and Pacific-Arctic. Finally, no genetic differentiation was observed between populations in the Kara-Laptev and the Atlantic-Arctic regions. This was rather expected given that a unique *Tara* Oceans Station in the Atlantic-Arctic, TARA_175, appeared to harbour *Chaetoceros* populations in the present analysis and is located at the interface between the Kara-Laptev regions (Supplementary Figure S12).

### Examining the correlation of abiotic parameters with population structure

The above results show that, depending on the MAG considered, there are noticeable patterns of population structure among *Chaetoceros* populations. We then investigated the correlation of different environmental parameters and geographic distance with the genetic differentiation of the MAG populations. For this, we selected the MAGs that were present in at least three different stations or depths and with a variance superior to zero, namely MAGs ARC_116, ARC_217 and SOC_37, all of them having populations in the Arctic Ocean. Pairwise-*F*_ST_ values between the MAG populations were modelled depending on a range of environmental parameters and Euclidean distance by applying a linear mixed model (LMM), as described in Laso-Jadart et al. (2021), to perform a variance partitioning analysis. The fixed part of the unexplained variance was below 10% for the three analyses, and was therefore considered negligible. We further applied Mantel tests to verify these results. For ARC_116, most of the genomic variation was correlated with silicate concentrations (35%), followed by phosphate (14%) and nitrite concentrations (13%), but they were not validated by the Mantel tests (Supplementary Figure S14A-C). A small correlation with the geographic distance was noted (7%), which was validated by the Mantel test (Supplementary Figure S14D).

On the other hand, iron was the environmental parameter correlated the most (77%) with genomic differentiation of ARC_217, which was not significantly validated by a Mantel test (Supplementary Figure S15). Finally, it was phosphate (45%), nitrite (14%), silicate (13%) and temperature (11%) that were the most correlated with genomic differentiation of SOC_37 populations. Phosphate, nitrite and silicate were not validated by the Mantel tests but temperature was (Supplementary Figure S16A-D). The fact that some of the Mantel tests were not validated despite a strong correlation of one parameter in the variance partitioning analyses was expected given that most of our samples were small, particularly for ARC_217 and SOC_37. It is indeed evident that most of the data points for these two MAGs are fairly dispersed around the regression curve, with minor exceptions (Supplementary Figures S14-S16). Moreover, Mantel tests may sometimes give biased *p*-values given the autocorrelation of some environmental variables examined in ecology studies (Guillot and Rousset 2013; Diniz-Filho et al. 2013). Taking this into account, some Mantel tests nonetheless confirmed the correlation between abiotic parameters and genetic differentiation of *Chaetoceros* populations observed in the variance partitioning analyses. From this we conclude that micro-diversification appeared to occur in response to different environmental factors in at least some closely related *Chaetoceros* populations.

### Identification of *Chaetoceros* genes under selection

Given the correlation observed between abiotic parameters and the *Chaetoceros* population structure patterns, we then examined whether some genes were undergoing selection. This analysis was conducted on the MAGs presenting variants in at least 3 different populations with discriminant *F*_ST_ values, namely MAGs ARC_116, ARC_217 and SOC_37. The LK distribution of the respective loci followed the expected chi-square distribution (Supplementary Figure S17), indicating that the loci followed the neutral evolution model of a single species. We identified several strong candidates potentially under positive selection on the genome contigs, that is 28 loci for ARC_116, 1,116 loci for ARC_217 and 1,101 loci for SOC_37, representing 0.17% (28); 6.89% (802) and 6.60% (534) of the genome contigs, respectively (Supplementary Table S6A-C). Globally, an inspection of the B-allele frequencies (BAF) showed that loci under selection were derived principally from *Tara* Oceans stations 175, 188, 193 and 201 (all SUR) for ARC_116, while it was mainly *Tara* Oceans station 201 (SUR or DCM) for ARC_217 and stations 173 and 201 (both DCM) for SOC_37.

We examined the Gene Ontology (GO) terms generated during the Interproscan analysis to gain insights into the functional repertoire of the genes under selection. All three MAGs presented GO terms associated with cellular components, with ARC_116 displaying a more elevated proportion of genes associated with membrane domains (Supplementary Figure S18). Regarding molecular functions, the three MAGs displayed an elevated proportion of genes associated with binding and catalytic activities, and ARC_217 showed the most diverse GO terms in this category. The GOs associated with biological processes were mostly represented by general cellular and metabolic processes for all three genomes, followed by cellular organisation or biogenesis as well as localization.

To focus our analysis on describing the potential functions associated with the genes under selection, we selected the SNVs located within a gene sequence with an assigned PFAM domain. A total of 31 PFAM domains were found in the genes harbouring the loci under selection for ARC_116, while we identified 697 for ARC_217 and 805 for SOC_37. Among the PFAMs associated with ARC_116 loci 54.84% (17) presented an associated GO term while the percentage was lower for ARC_217 (46.92%) and SOC_37 (40.75%). Most of the outlier loci were associated with coding or untranslated regions (UTRs), with a minor contribution of loci within intronic regions (Supplementary Figure S19). Some variants were responsible for loss of function events, such as stop codon gain and frame-shift mutations. We subsequently searched for domain functions potentially linked to the environmental parameters possibly driving the micro-diversification patterns.

Strikingly, all variants under selection within the ARC_116 populations were completely absent from station TARA_189 (DCM). Most of the non-synonymous loci displayed domains associated with kinases, oxidoreductases and transferases. Among the loci under selection were domains involved in redox balance, such as missense or 3’ UTR variants in genes harbouring glutaredoxin (PF00462, PF13417) and cytochrome *c* oxidase domains (PF02683), all mostly fixed in surface populations of stations TARA_175, TARA_193 and TARA_201 (Supplementary Table S6D). Other loci included synonymous, missense and 3’ UTR variants in domains involved in intracellular transport (e.g., PF04811 and PF08318). Both these domains appeared associated with endoplasmic reticulum to Golgi transport and their variants were almost fixed at the surface of stations TARA_175, TARA_188, TARA_193 and TARA_201.

Conversely, both ARC_217 and SOC_37 showed loci under selection for potential chlorophyll-binding and CobW proteins. The former are found in light-harvesting complexes of the photosynthetic apparatus while the latter form a large family of metal chaperones associated with metal homeostasis processes either with zinc, iron or cobalt molecules (Haas et al. 2009; Hsieh et al. 2013). Interestingly, both these functions may have a link with iron availability status, as phytoplankton can cope with iron limitation through remodelling of light-harvesting complexes, while some CobW proteins may exert iron-responsive patterns (Behrenfeld and Milligan 2013; Kotabova et al. 2021). Comparing the frequency of these variants among the two genomes, both ARC_217 and SOC_37 populations were found at stations TARA_201 (Arctic archipelago) and TARA_173 (Kara-Laptev) and showed elevated frequency of this variant at the former, whereas those at station TARA_173 exhibited lower frequency (Supplementary Table S6E-F). Looking at the environmental metadata of these stations using the PANGAEA database (Ardyna et al. 2017; Guidi, Morin, et al. 2017; Guidi, Picheral, et al. 2017), we observed that station TARA_173 was characterised by higher iron and nitrate levels but was lower in phosphate (Fig. 6E, Supplementary Table S5). ARC_217 populations were also found at station TARA_205 (SUR), which displayed the lowest iron concentration (51% and 38% lower than the ones of stations TARA_173 and TARA_201 (both SUR), respectively, Fig. 6E, Supplementary Table S5), where the CobW variant was completely fixed and the chlorophyll-binding domain was completely absent. Moreover, we identified a synonymous variant of ARC_217 located in gene TARA_ARC_108_MAG_00217_000000002161.2.2 within a flavodoxin domain (PF00258). Flavodoxin proteins are known to over-accumulate in iron-limited conditions over their iron-containing counterpart ferredoxin (La Roche et al. 1996). This SNV was almost fixed at the surface of stations TARA_201 and TARA_205 (Supplementary Table S6E). A variant within an iron-sulfur cluster associated domain (PF02657) involved in redox and regulation of gene expression processes (Johnson 1998) was also found almost fixed in TARA_173 (DCM) and TARA_201 (SUR) SOC_37 populations (Supplementary Table S6F).

**Figure 5.**
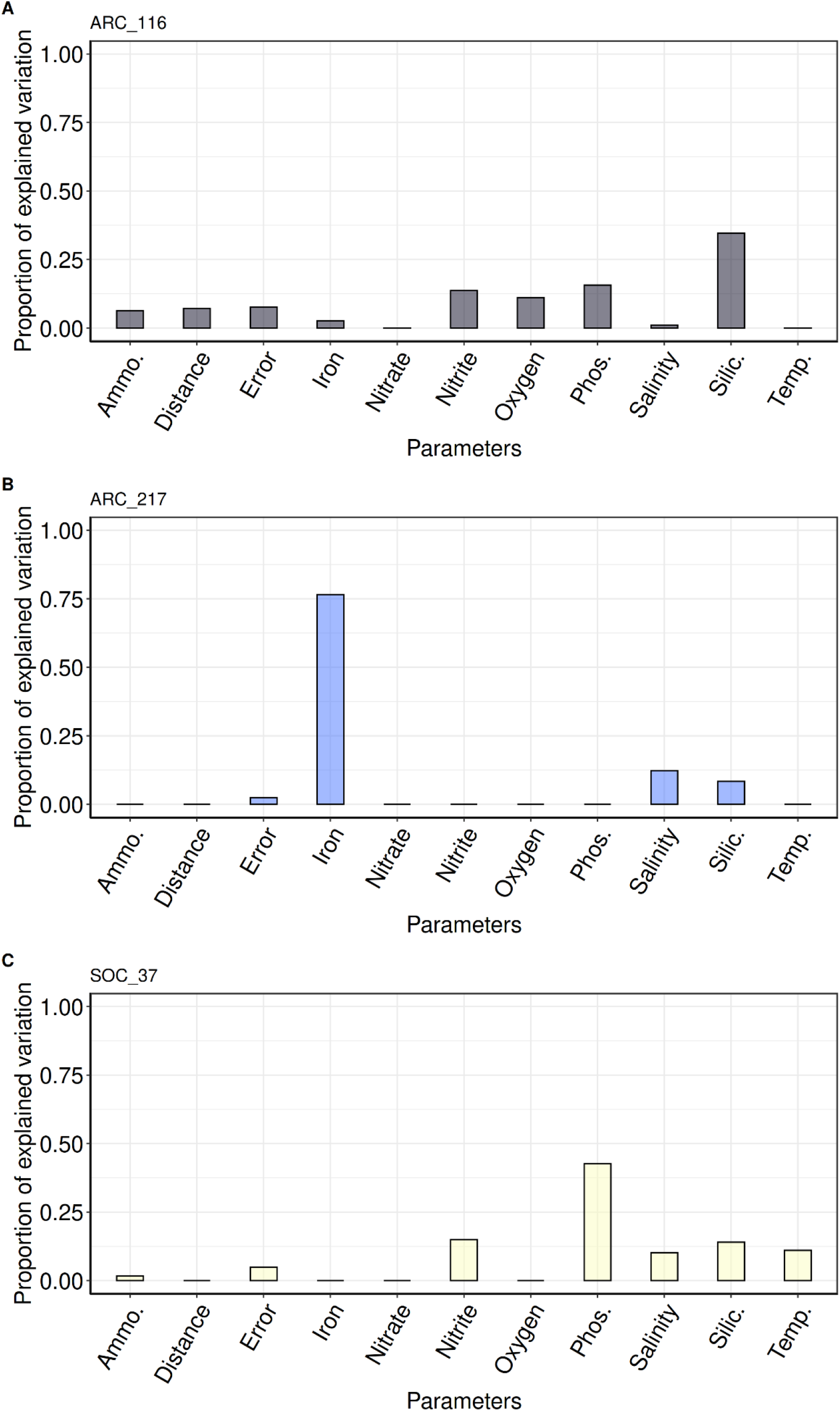
Correlation between environmental parameters variation and *Chaetoceros* genomic differentiation. Barplots of variance partitioning analysis results for (A) ARC_116, (B) ARC_217 and (C) SOC_37. Ammo.: ammonium; Phos.: phosphate; Silic.: silicate; Temp.: temperature.

**Figure 6.**
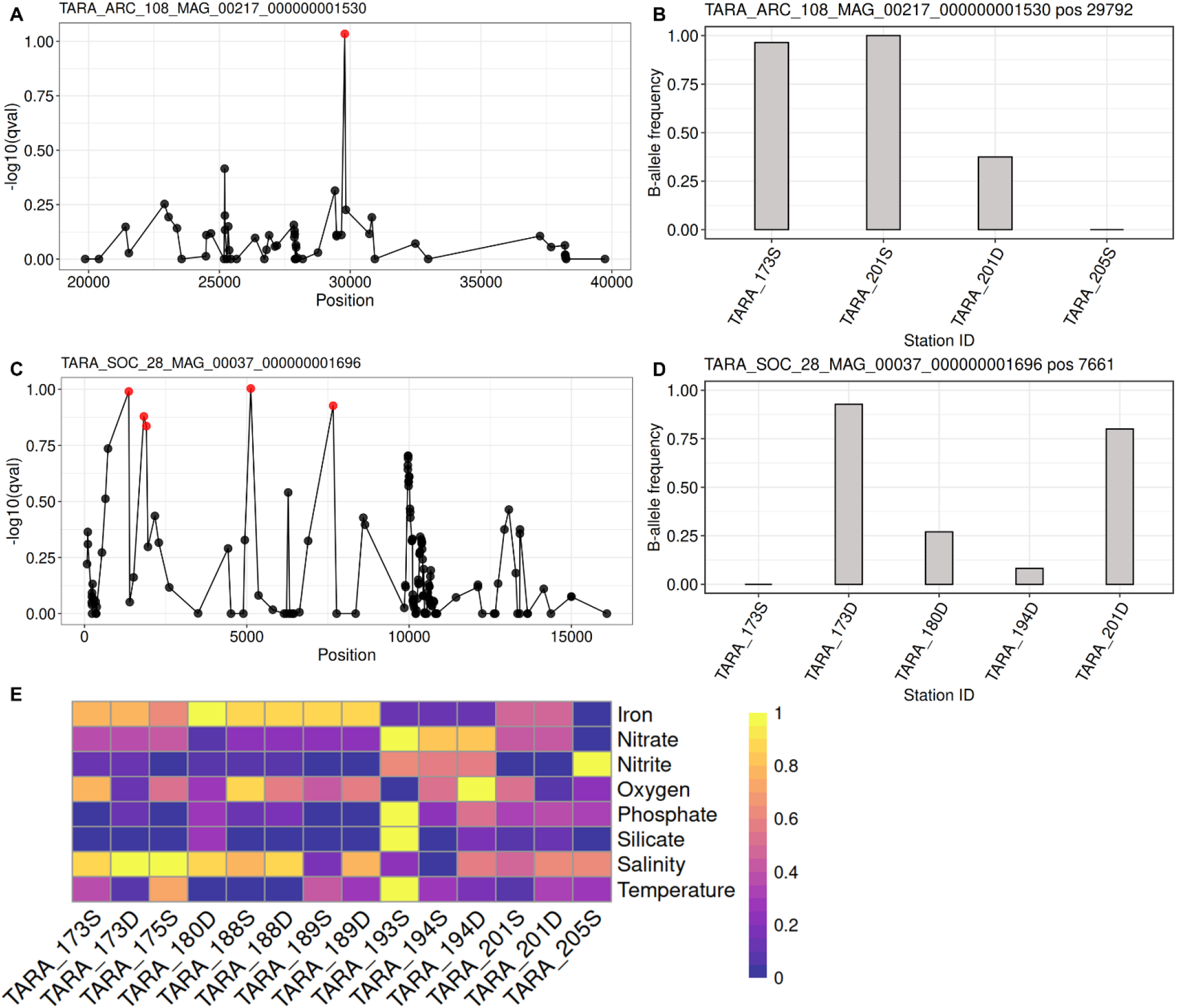
Selection of variants in Arctic *Chaetoceros* populations. (A) and (C) represent Manhattan plots in a 10 kb window around the variants of interest, shown for ARC_217 (ISIP) and SOC_37 (spermidine/spermine synthase). Red dots correspond to SNVs considered under selection (*q*-value < 0.15). (B) and (D) represent the barplots of the B-allele frequency (BAF) for the respective loci of interest depending on the population considered. (E) Range-transformed heatmap of abiotic parameters for the *Tara* Oceans stations in the Arctic Ocean where the *Chaetoceros* MAGs are present (see Supplementary Table S5 for raw values). The brightest yellow colour represents the most elevated values for any given parameter in the dataset, while the darkest purple indicates the lowest. D: deep-chlorophyll maximum; S: surface.

Most notably, one variant of ARC_217 appeared to be located in a gene encoding a low iron-inducible periplasmic domain (PF07692). This SNV corresponded to a synonymous mutation located in the coding region of gene TARA_ARC_108_MAG_00217_000000001530.105.1. An examination of the Manhattan plot of this SNV revealed few drafted variants around the loci under selection, indicative of a potential soft selective sweep, and the BAF showed that selection of this variant was occurring on *Chaetoceros* populations from stations with higher iron concentrations (TARA_173 and TARA_201; both SUR) (Fig. 6E, Supplementary Table S5), while it was completely absent from TARA_205 (SUR) (Fig. 6A-B). Next, we searched for potential homologue candidates in *Phaeodactylum tricornutum* by aligning the corresponding ARC_217 protein on the Phatr3 proteome (Rastogi et al. 2018) using BLASTp. We identified protein B7FYL2 (https://bioinformatics.psb.ugent.be/plaza/versions/plaza_diatoms_01/genes/view/ptri151180) which was previously identified as an iron starvation-induced protein (ISIP) ISIP2a (Bowler et al. 2008), and showed an e-value of 1e-13. The corresponding gene located within a genomic region marked by histone post-translational modification (PTM) in H3K4me2, a mark suggested to associate with expressed genes in *P. tricornutum* (Veluchamy et al. 2015). The mark was reduced in nitrate-limited conditions compared to repletion. We further compared homologue sequences among the 8 *Chaetoceros* transcriptomes from Marine Microbial Eukaryote Transcriptome Sequencing Project (MMETSP) that were used for the phylogeny reconstruction (see Results) and retrieved 16 candidate sequences from 7 transcriptomes (see Supplementary data file S1, Supplementary material online). Aligning these sequences to the reference gene allowed us to identify significant identity at the SNV position for 7 (44%) sequences, including 2 sequences from *C. dichaeta* that displayed the same SNV (C>T) as the loci under selection (Supplementary Figure S20, Supplementary data file S2, Supplementary material online). Although the SNV observed in the ARC_217 gene sequence was predicted to induce a synonymous mutation, we searched whether the nucleotide substitution modified the predicted RNA secondary structure through an analysis conducted on the LinearFold and RNAfold web servers (Gruber et al. 2008; Huang et al. 2019), as silent mutations may impact haplotypes as for instance changes in RNA secondary structure (Sauna et al. 2007). No clear change of RNA secondary structure was predicted by LinearFold (Supplementary Figure S21A-B). RNAfold outputs showed that the reference sequence displayed an ensemble diversity (i.e., an average base-pair distance between all the structures in the thermodynamic ensemble) of 1514.62, while the mutated sequence showed a value of 1486.24. Slight differences of free energy minimisation and centroid (structure with the minimal total base-pair distance to all structures in the ensemble) structures (Supplementary Figure S21C-D) were observed. These results suggested a possible minor impact of the mutation on the RNA folding structure.

Next, an examination of SOC_37 loci under selection revealed genes encoding functions associated with phosphate metabolism, such as a putative cytosolic domain of 10TM phosphate transporter (PF14703), protein and histidine phosphatase domains (PF00481 and PF00300) (Supplementary Table S6F). In accordance with its enrichment in RNA polymerase Rpb1 C-terminal domain (see Results), many genes under selection harboured PFAM domains associated with RNA, as for instance an elongation factor of RNA pol II, RNA recognition motifs and binding domains, in addition to a reverse transcriptase involved in transposable element activity, most of them almost fixed in station 201 (DCM) while absent from station TARA_173 (SUR) and under selection at the DCM of stations TARA_173, TARA_180 and TARA_194 (Supplementary Table S6F). Other functions notably included domains potentially involved in methyl transfers and epigenetic mechanisms regulating gene expression (e.g., PF00850, PF00856 and PF08123), all encoded by genes under selection in TARA_173 and TARA_201 (DCM). Most remarkably, loci under selection included two variants potentially involved in polyamine biosynthetic processes. One SNV was located in the UTR of carbamoyl phosphatase domains (PF00988 and PF02786) and the other in the coding region of a spermine/spermidine synthase domain (PF01564) in gene TARA_SOC_28_MAG_00037_000000001696.30.1, causing a non-synonymous mutation (p.Met1063Leu). Polyamines such as spermine and spermidine are involved in frustule formation through the production of long-chain polyamines (Kröger et al. 2000) using carbamoyl phosphate synthase (Armbrust et al. 2004). Inspection of the Manhattan plot of the polyamine synthase variant suggested a potential hard selective sweep signature, and the BAF showed that it was almost fixed at the DCM of stations TARA_173 and TARA_201 (Fig. 6C-D) which were characterised by the lowest oxygen concentrations in our analysis (Fig. 6E, Supplementary Table S5). The carbamoyl phosphate variant showed similar patterns and appeared more frequent in stations with less nitrate and phosphate (Supplementary Tables S5 and S6F). A sequence similarity search for a polyamine synthase homologue in the Phatr3 proteome allowed us to identify the protein B7FPJ4 (https://bioinformatics.psb.ugent.be/plaza/versions/plaza_diatoms_01/genes/view/ptri221670) involved in spermidine biosynthesis (Bowler et al. 2008), with an e-value of 3e-163. The gene coding this protein exhibited significant changes of expression in nitrate depletion compared to repletion (- ~0.5 fold change; Levitan et al. (2015)) and in phosphate depletion compared to repletion (~2.9 fold change; Cruz de Carvalho et al. (2016)). Moreover, this gene located in region marked by H3K9me2 and H3K4me2 PTMs in *P. tricornutum* (Veluchamy et al. 2015). Searching among the 8 *Chaetoceros* transcriptomes from MMETSP yielded 17 candidate sequences from 3 transcriptomes (see Supplementary data file S3, Supplementary material online). Among them, 16 (~94%) sequences showed significant identity at the SNV position with 1 sequence from *C. debilis* displaying the same SNV (A>C) as the loci under selection (Supplementary Figure S22, Supplementary data file S4, Supplementary material online). Predictions using LinearFold confirmed clear changes of RNA secondary structure between reference and mutated gene (Supplementary Figure S23A-B). Moreover, RNAfold predictions agreed with this pattern as they indicated an ensemble diversity of 957.66 and 1039.87 for the reference and mutated sequences, respectively, along with significant modifications of free energy minimisation and centroid structure (Supplementary Figure S23C-D). Overall, these findings illustrate the process by which a single mutation may have a direct effect on the electrochemical properties of RNA structures, as well as potentially impacting the biochemical kinetics of the protein.

## Discussion

Our understanding of both the ecology and biological functions of marine algae has progressed considerably with the help of molecular-based methods, generating an increasing number of genomes now reaching more than several hundred (Hanschen and Starkenburg 2020; Grigoriev et al. 2020). Nonetheless, one drawback of laboratory-generated genomes is their potential lack of representativeness of species that thrive in the environment. As an example, it has been demonstrated that *P. tricornutum* cultures artificially selected individuals in the population, thus reducing the overall diversity of their genetic pool leading to genetic convergence of the strains; with nutrient-replete conditions favouring somatic mutations leading to the loss of function of some genes that are potentially important in highly fluctuating environments (Helliwell et al. 2015; Rastogi et al. 2020). Collectively, these factors may constitute a limit to the understanding of metabolism and locally relevant genomic functions. Moreover, interrogating *Tara* Oceans metagenomes with the *Fragilariopsis cylindrus* CCMP 1102 genome showed enough read coverage for only one sampling station, further exemplifying the divergence between laboratory strains and natural populations (Bulankova et al. 2021). While cultivating strains in the lab is a necessary first step to gaining insights into their fundamental ecology, accessing the genomes of organisms directly from their environment is key to fully understanding their role within natural communities and their responses to environmental fluctuations. In this context, genome-resolved metagenomics, which consists in the mapping of environmental metagenomic reads on a reference genome, represents a powerful tool to access the diversity and distribution of organisms in their native environment without relying on taxonomic markers that may be too conservative to unveil the amount of diversity, a fact that appears in particular to be the case for unicellular organisms (Piganeau et al. 2011). However, as this field is in its infancy, specific attention must be paid during the binning of metagenomes to prevent chimeric assembly (Nelson et al. 2020). Metagenome-assembled genomes must furthermore include information about assembly quality, level of contamination and completeness, to enable robust comparisons between studies (Bowers et al. 2017). The exploration of marine phytoplankton population genomics using metagenomes has only just begun, with a few pioneering studies focused on the Mamiellales genera *Bathycoccus*, *Micromonas* and *Ostreococcus* (Leconte et al. 2020; Leconte et al. 2021). In line with these, the present study aimed to generate a portrait of the diversity landscape among natural *Chaetoceros* populations and to bridge the gap between diatom genomes, physiological responses and population genomics by teasing apart the correlation of geographical distance and environmental factors with population structure. Going from not only one but eleven metagenomes to gene selection using genome-resolved metagenomics, this work is to our knowledge the first to assess population-scale diversity among *Chaetoceros* genomes directly reconstructed from the environment.

### *Chaetoceros* metagenome-assembled genomes from *Tara* Oceans

The genomes we considered for the present study have been generated from the *Tara* Oceans metagenomic reads and displayed the same magnitude in size, number of protein-coding genes and G+C content as the newly published *Chaetoceros tenuissimus* genome (Hongo et al. 2021), pointing out the accuracy of the MAG reconstruction methods. Several studies have investigated the link between genome size compared to cell morphology and metabolism and have found that both cell size and growth rate are, respectively, proportional and inversely proportional to genome size (Williams 1964; Holm-Hansen 1969; Shuter et al. 1983; Veldhuis et al. 1997; Cavalier-Smith 2005; Von Dassow et al. 2008). We observed contrasted differences in genome sizes for the MAGs ARC_232, ARC_267, MED_399 and PSW_256, which all exhibited the same level of G+C% (the lowest being around 39%), displayed an enrichment of the level of D, E, K and N amino-acids and a depletion in C, D and A residues, and seemed to belong to the *C. affinis* subclade, indicating that they potentially belong to the same species. These genome size discrepancies suggest potential contrasted growth rates and cell sizes for closely related *Chaetoceros* species. Koester et al. (2010) noted a two-fold genome size difference between cryptic but geographically separated populations of the diatom *Ditylum brightwellii*, accompanied by a difference in growth rate, suggesting that whole-genome duplication events may constitute important drivers of genetic diversification in diatoms. Here, both ARC_232 and ARC_267 populations were identified in the Arctic but did not seem to co-occur in our analyses, while MED_399 and PSW_256 were found to be restricted to the Mediterranean and Pacific Oceans, respectively. It is possible that duplication and/or transposition events potentially linked to stress, as was observed in *P. tricornutum* (Maumus et al. 2009), gave rise to diverged subpopulations that dispersed and were finally genetically separated following allopatric speciation.

### Insights into biogeographical patterns of the genus *Chaetoceros*

We identified each of the MAGs in a relatively small number of samples with an uneven distribution, suggesting potential habitat specialists, with the notable exception of ARC_116, the only MAG that was distributed in samples spanning globally across the Arctic Ocean, indicating a potential pan-Arctic species. Other studies, based on the *Tara* Oceans and Ocean Sampling Day sample datasets, have already provided a thorough pattern of *Chaetoceros* distribution at global scale in the oceans (Malviya et al. 2016; De Luca, Kooistra, et al. 2019). These have shown a prevalence of *Chaetoceros* in the Arctic Ocean, with discrepancies depending on the species considered. For instance, metabarcoding analyses showed that *Chaetoceros neogracile* was restricted to the northern hemisphere (De Luca, Kooistra, et al. 2019), which consistently matches the distribution of ARC_116 and its taxonomic closeness to *C. neogracile* RCC1993. *C. dichaeta* has been retrieved near Alaska and the Antarctic peninsula, a distribution that appears in line with that of ARC_189 and with the geographic closeness of PSE_253 in South America. The same study indicated that *C. affinis* was present in the Mediterranean as well as in the Atlantic Ocean and North Sea. We found 4 MAGs (ARC_232, ARC_267, MED_399 and PSW_256) that exhibited taxonomic closeness to *C. affinis*. Only one of them was found in the Mediterranean Sea while the three others were retrieved from the Pacific and Arctic Oceans, suggesting potential new niches for this species. *C. debilis*, of which our MAG SOC_60 was also found to be very close, was retrieved in different localities: in European coastal waters and in the Arctic Ocean for the northern hemisphere, as well as in the Indian and Southern Oceans, in agreement with the distribution of this MAG. *Chaetoceros* has been viewed as a local opportunistic genus (Barton et al. 2010; Smodlaka Tanković et al. 2018) but the various species we describe here appeared to evolve in sympatry with up to three MAG populations in the same sampling stations. A temporal survey of these sampling points could help reveal whether the populations from different species are sympatric on a regular manner or if some competition mechanisms or niche exclusions are observable. Of note is the observation that none of the MAGs belonging to the same subclade were found at the same locations, with the exception of ARC_217 and SOC_37. In general, we found most of our *Chaetoceros* populations in the Arctic Ocean, which may be due to their lower sequence coverage in tropical and subtropical waters, as these species are likely to be among dominant phytoplankton in the Arctic (Sommeria-Klein et al. 2021). It is moreover evident that other localisations harbour *Chaetoceros* populations, such as for instance in the Southern Ocean where diatoms dominate photosynthetic protist assemblages (Malviya et al. 2016; Sommeria-Klein et al. 2021). In the same vein, associations between *Chaetoceros* and tintinnid ciliates have been observed in the Pacific Ocean and Caribbean Sea (Gómez 2007; Gómez 2020), which were only partially sampled during the *Tara* Oceans expeditions (Malviya et al. 2016). Future oceanographic campaigns should help reveal the distribution of *Chaetoceros* populations and the extent of their genomic variability.

### Genetic differentiation among closely related *Chaetoceros* populations is correlated with different environmental variables

We investigated global patterns of population structure and genetic differentiation in *Chaetoceros*, a cosmopolitan diatom found in every major oceanic province, and one of the most diverse. By leveraging metagenomes reconstructed through the *Tara* Oceans expeditions, we drew a comprehensive landscape of the genetic diversity among different populations of this genus and were able to address their level of gene flow through an analysis of their population structure. The levels of micro-diversity observed here, ranging from 0.63% to a maximum of 2.34%, are in line with previous analyses conducted on natural populations of the diatom *Fragilariopsis cylindrus* from *Tara* Oceans station 86, which displayed ~2% SNV density (Bulankova et al. 2021). The observation of elevated genetic differences among populations from the same species means that despite their high potential dispersal, *Chaetoceros* diatoms can express significant levels of divergence. Moreover, our variance partitioning analyses revealed that the genetic differentiation between *Chaetoceros* populations was correlated with a combination of different abiotic factors, with only a minor correlation with geographic distance. This is in agreement with previous studies analysing population diversity using microsatellite markers, such as Härnström et al. (2011) on *Skeletonema marinoi* and Whittaker and Rynearson (2017) on *Thalassiosira rotula*, and contradicts the results found by Casteleyn et al. (2010) for *Pseudo-nitzschia pungens*. It must be noted that the former two are homothallic centric diatoms while *P. pungens* is a heterothallic pennate diatom. Therefore, the historical assumption that geographic distance is the parameter conditioning most microbial genetic diversity appears conflicting in diatoms.

Among the most notable nutrients that regulate diatom populations are nitrate (Moore et al. 2004), iron (Boyd et al. 2007; Caputi et al. 2019), phosphate (Egge 1998; Cruz de Carvalho and Bowler 2020), silicon (Martin-Jezequel et al. 2000; Yool and Tyrrell 2003) and cobalamin (i.e., vitamin B12) (Bertrand et al. 2012; Ellis et al. 2017), although environmental controls of diatom populations vary locally due to their cosmopolitan nature. To our knowledge, the closest study to the present one is that of Whittaker and Rynearson (2017), where the authors investigated the correlation of abiotic parameters and geographic distance with *Thalassiosira rotula* population structure and revealed a correlation with temperature. By contrast, in the present analysis we rather show a correlation of genetic differentiation with phosphate, silicate and iron in *Chaetoceros* species, with only a minor (~10%) correlation between temperature and genetic differentiation in SOC_37 populations. It should nonetheless be noted that the MAGs were found in samples displaying a narrow range of temperature between 1 and 4 °C. Therefore, the patterns of environmental control on global genomic diversity are consistent with expectations from the literature.

Significant population structure was observed among the *Chaetoceros* MAGs, but with relatively moderate between-region differences in the Arctic, as was observed for zooplankton (Laso-Jadart et al. 2021, p.), with *F*_ST_ levels reaching up to ≥ 0.2. These high levels of genetic differentiation appear approximately close to those described in different *P. tricornutum* accessions (pairwise *F*_ST_ ~0.2-0.4), represented for the most part by strains that have been maintained in culture collections for decades (Rastogi et al. 2020). In particular, ongoing speciation of the ARC_116 population located at station TARA_194, particularly at the DCM, might be a reason explaining why we observed a dramatic number of SNVs, leading us to exclude this station in order to perform a more conservative study when identifying genes under selection. Indeed, this indicates unequal gene flow among the populations and suggests a metapopulation structure consisting of populations of populations, as has been described for the diatom *D. brightwellii* (Rynearson et al. 2009). This difference in genetic structure of the ARC_116 population at station TARA_194 was not observed in other populations present at this particular station, such as for SOC_37. Overall, all three MAGs ARC_116, ARC_217 and SOC_37 appeared closely related in our phylogenetic analyses but nonetheless showed elevated numbers of MAG-specific orthogroups, and their genetic differentiation was correlated with different sets of abiotic parameters. Taken together, these results emphasize that even with the same local environmental conditions, populations of closely related diatoms from the same genus do not display identical gene flow patterns, emphasising their enormous genetic diversity as well as significant adaptive potential.

### Functional overview of natural selection among *Chaetoceros* populations

We were able to identify genes under selection between the different *Chaetoceros* populations and tried to assess their respective functions. Previous studies have tried to investigate the gene functions that are essential for diatom survival. Among these are functions associated with light perception and energy dissipation, such as for instance phytochromes involved in red/far-red light sensing (Fortunato et al. 2016) and light-harvesting complex stress-related proteins (LHCX1) that modulate light responses (Bailleul et al. 2010). Other important functions include metabolic plasticity and response to nutrient fluctuations. As an example, ornithine-urea cycle proteins mediate rapid responses to nitrogen variability (Allen et al. 2011). Another remarkable characteristic of diatoms is their ability to respond to iron fluctuations, as they are among communities most strongly linked to the concentration patterns of this micronutrient (Caputi et al. 2019). Indeed, diatoms exhibit a diverse range of iron uptake mechanisms, involving siderophores (Kazamia et al. 2018), phytotransferrins (Allen et al. 2008; Morrissey et al. 2015) and ferric reductases (Gao et al. 2021). They exhibit differential mechanisms of iron storage, with *Pseudo-nitzschia* using ferritin and members of the *Chaetoceros* and *Thalassiosira* genera are believed to be able to store iron in their vacuole (Lampe et al. 2018). While we did not observe selection of functions related to light acquisition in our analyses, clear positive selection patterns of genes involved in iron response were retrieved, for the most part in ARC_217 populations. These included a gene encoding an iron starvation-induced protein (ISIP), which exhibited contrasted frequency that followed iron concentration patterns, the variant being more frequent in stations with more elevated iron concentrations. We found this gene to be a homologue of *P. tricornutum* ISIP2a, which encodes a protein involved in concentrating ferric iron at the cell surface (Morrissey et al. 2015), with a function equivalent to human transferrin, hence its name “phytotransferrin” (McQuaid et al. 2018). This protein has been proposed to constitute an ecological marker of iron starvation in diatoms (Marchetti et al. 2017) as it is strongly up-regulated by this condition. Previous analyses in *P. tricornutum* cells deficient in ISIP2a showed reduced iron uptake capabilities (Morrissey et al. 2015; Kazamia et al. 2018). While our results suggested a slight effect of the mutation on its RNA secondary structure, we identified the same SNV in sequences homologous to this gene in *Chaetoceros* transcriptomes. It therefore appears plausible that relaxed selective pressure on the gene in an environment more iron-replete could have led to the observed mutation.

Additionally, we noted the positive selection of genes encoding a carbamoyl phosphate synthase and spermine/spermidine synthase in SOC_37 populations, with the latter showing a potential impact on RNA secondary structure, along with other genes linked with phosphate metabolism. Carbamoyl phosphate synthase is thought to catalyse the first urea cycle step, a process that generates polyamine precursors (Armbrust et al. 2004). Polyamines such as spermine and spermidine are nitrogenous compounds involved in frustule formation through their interaction with the heavily phosphorylated silaffin phosphoproteins (Kröger and Sumper 1998; Poulsen et al. 2003). In consequence, diatom frustule formation relies on both nitrogen and phosphorus. We identified a polyamine synthase homologue in *P. tricornutum* displaying significant modulation of its expression in response to nitrate and phosphate availability levels, as well as homologue bearing the same mutation in a *Chaetoceros* transcriptome. Polyamine biosynthetic processes have been linked to diatom physiological responses to nitrogen, salinity and temperature (Scoccianti et al. 1995; Liu et al. 2016; Gleich et al. 2020) and many polar diatoms show increased silicate content under iron-limited conditions, which can result from either increased silicate accumulation or lower accumulation of nitrate depending on the species considered (Timmermans et al. 2004; Hoffmann et al. 2007). Despite this, the putative link between abiotic parameter variations across stations and the variant frequency patterns observed remains unclear. In this study we noted that several genes harbouring SNVs displayed homologues associated to PTMs in *P. tricornutum* (Veluchamy et al. 2015). Changes in the amount of cytosine residues in genes (e.g., A>C) could potentially affect gene expression through the increase or decrease of methylation sites available, especially when in CpG islands context. Overall, future studies involving engineered knock-out mutants of the genes containing the SNVs and including different sets of abiotic parameters should help gain insights into their respective impact on gene expression patterns and whole-cell fitness.

### Conclusion

*Chaetoceros* is the most widespread and connected diatom genus, making it a key component of plankton communities, and has been identified as a genus vulnerable to projected climate change. Here, we have shown significant correlations between nutrient availability and genetic differentiation among *Chaetoceros* populations, with a potential impact on their growth strategies. As climate change is expected to influence water stratification, acidification as well as nutrient availability, it appears more than likely that predicted environmental changes in the Arctic will influence *Chaetoceros* distributions and its gene pool. The present study positions itself as an extension of previous work realised on plankton population genomics, and is to our knowledge the first of its kind dealing with micro-diversification patterns of multiple metagenome-assembled genomes from a non-model diatom genus. Finally, this work highlights the necessity to perform repeated sampling over time to be able to test whether separated *Chaetoceros* populations can evolve in sympatry as well as the effect of seasonality and local environmental fluctuations on gene selection.

## Materials and Methods

### Genomic resources

Eleven reconstructed and manually curated metagenome-assembled genomes (MAGs), generated from *Tara* Oceans metagenomic reads (Delmont et al. 2022); available at https://www.genoscope.cns.fr/tara/#SMAGs) and belonging to the genus *Chaetoceros* were considered in the present study. For clarity and readability, the original MAG IDs were shortened and the corresponding information is available in Supplementary Table S1. Information corresponding to the genome size and number of genes were extracted from Delmont et al. (2022). All these genomes received a former geographical assignment based on their read recruitment in the *Tara* Oceans sampling stations after mapping each MAG onto the *Tara* Oceans metagenomic dataset. The estimation of genome completion was performed by retrieving the Benchmarking Universal Single-Copy Orthologs (BUSCOs) (Simão et al. 2015) using the DNA contigs of the MAGs and control genomes as input for BUSCO v2.0.0 with the eukaryota_odb10 library. Gene length was estimated using SAMtools (Li et al. 2009) ‘faidx’ on the genome FASTA files. Percentage of G+C in the genome codons was calculated with the COUSIN online tool (Bourret et al. 2019). Reference diatom genomes for *Phaeodactylum tricornutum* CCAP 1055 and *Thalassiosira pseudonana* CCMP 1335 were also considered to provide a comparison with the *Chaetoceros* MAGs and were both retrieved from the Joint Genome Institute (Armbrust et al. 2004; Bowler et al. 2008). Information about their completion level, gene length and percentage of GC were obtained with the aforementioned methods.

### Average nucleotide identity and average amino-acid identity

Average nucleotide identity (ANI) between the MAGs was calculated using FastANI (Jain et al. 2018) on the whole genomes with the --ql and --rl parameters and --minFraction 0.05 to retrieve all the identity percentages even for highly divergent MAGs. In a similar manner, average amino-acid identity (AAI) was estimated on the whole predicted proteomes using the online AAI calculator provided by the Konstantinidis lab (http://enve-omics.ce.gatech.edu/aai/) (Rodriguez-R and Konstantinidis 2014).

### Phylogeny and identification of orthogroups

To investigate the taxonomic relatedness of the MAGs, we considered the BUSCO genes identified from estimation of genome completion (part 3.1). Using the BUSCO IDs for which at least 8 out of the 11 MAGs (~70%) presented a sequence (83 over 255 eukaryotic BUSCOs), the BUSCO gene clusters translated into proteins were aligned with MAFFT (Katoh et al. 2002) in automatic mode, followed by a manual cleaning of each of the alignments by removing N-terminal and C-terminal residues displaying less than 70% conservation. The alignments were then trimmed using trimAl (Capella-Gutierrez et al. 2009) with the parameter -gt 0.5, followed by another alignment step with MAFFT. A total of 83 gene clusters were retained. To identify potential contaminants, consensus guide trees were generated for each of the approved alignments using RAxML (Stamatakis et al. 2005) with the parameters -m PROTGAMMAJTT for the substitution model, -N 100 bootstrap replicates and randomly defined numbers between 1 and 99,999 for the parameters -x and -p. To evaluate the MAGs relatedness with respect to other taxa, the same approach was conducted based on a concatenation tree with 23 supplementary taxa sampled across the eukaryotic tree of life (34 total taxa), including 8 *Chaetoceros* transcriptomes from MMETSP deposited in the European Nucleotide Archive converted into protein sequences, for the same 83 single-copy nuclear genes. The cleaned alignments were then concatenated and a final ML tree was built with RAxML 8.2.12 (100 bootstraps) and the final figure exported in iTOL (Letunic and Bork 2021). Protein sequences from the MAGs (available at https://www.genoscope.cns.fr/tara/#SMAGs) were used as input for the OrthoFinder (Emms and Kelly 2015) software to identify the orthogroups.

### Comparative analysis of the amino acid composition and PFAMs of the MAGs

The amino acid composition of the genomes was computed by analysing the FASTA files with the ‘protr’ package in R (Xiao et al. 2015) and plotting their corresponding frequencies. The amino acid composition of each MAG was then normalised by the respective amino acid global mean. Protein FAMilies (PFAMs) domains were inferred by searching in a local installation of Interproscan (Jones et al. 2014). Raw values are available in Supplementary Table S2.

### Genome-resolved metagenomics of *Chaetoceros* MAGs

To generate an estimate of *Chaetoceros* MAG abundance, we performed a mapping of the *Tara* Oceans metagenomic dataset on their contigs using BWA-MEM (Li and Durbin 2009) with default parameters, with an 80% identity filter and at least 4x mean vertical coverage, removed the duplicate reads, and stored the recruited reads as BAM files with SAMtools. The resulting mapping files were sorted using SAMtools ‘sort’ and the different size fractions were merged to increase the coverage, keeping the surface (SUR) and deep-chlorophyll maximum (DCM) depths separated in order to compare their corresponding local populations when possible. Read identity was extracted using a custom perl script using SAMtools ‘view’, and plotted in RStudio. We performed a final step of read filtration on their identity by extracting the read names with at least 97% identity in RStudio, then retrieving them using the Picard toolkit (https://broadinstitute.github.io/picard/) on the indexed BAM files with the option ‘FilterSamReads’ in ‘lenient’ mode. In order to ensure that the recruited reads belonged to our genomes of interest, we generated plots of the genomic coverage of the reads with Bedtools ‘genomecov’ (Quinlan and Hall 2010) and performed a visual inspection to detect bimodal trends. We excluded the reads from sampling stations that presented insufficient or bimodal coverage distribution (see Supplementary Fig. S3 where we report an example of discarded samples based on read coverage), and the ones with a coverage breadth estimated with SAMtools ‘depth’ inferior to 80. These read filtration and sample inspection steps were critical as some non-specific read recruitment may happen (due for instance to the stability of 18S rRNA gene and to hypervariable genomic regions). This resulted in a total of 20 *Tara* Oceans Stations and/or depths investigated. The relative abundance of the MAGs in each sample was estimated as the number of mapped reads, obtained with SAMtools ‘flagstat’, normalised by the total number of reads per sample (available in Supplementary Table S4 of Delmont et al. (2022)). We then computed the relative proportion of *Chaetoceros* reads per station and generated the corresponding maps with RStudio 4.0.1.

### Population genomics analyses

Genomic variants of the *Chaetoceros* populations associated to different stations and/or depths were called with BCFtools 1.13.25 (Li 2011) to generate a VCF file (mpileup of the files with at least 97% identity). Variant annotation of the VCF files was conducted using SnpEff (Cingolani et al. 2012), converting GFF files into GTF 2.2 format with AGAT (Dainat et al. 2022). The number, position and type of variants as well as their respective effects were plotted on RStudio.

To calculate the genetic distance between the MAG populations, the genetic variants were identified using SAMtools mpileup -B (multiple BAM files per MAG) and merged into one file. The Popoolation2 tool (Kofler et al. 2011) was then applied on the MAGs present in at least two stations (ARC 116, ARC 217, ARC 232, SOC 37) to generate a synchronised (.sync) file from the merged mpileup with the following parameters: --fastq-type sanger --min-qual 20. The .fst files were subsequently computed with the parameters --suppress-noninformative --min-count 2 --min-coverage 4 --max-coverage 200 --min-covered-fraction 1 --window-size 1 --step-size 1 --pool-size 500. The FST metrics were computed from the allele frequencies (not the allele counts) using the equation in (Hartl and Clark 2007). Allelic frequencies were computed with PoPoolation2 with the parameters --min-count 2 --min-coverage 4 --max-coverage 200. To ensure that these alleles belonged to our query genomes, the global population-wide F-statistic was computed for each MAG of interest and its distribution plotted and inspected for unimodality. Median pairwise-*F*_ST_ values were considered as a proxy for genomic differentiation between the respective MAG populations. In addition, the LK statistics (Lewontin and Krakauer 1973) were computed and compared with the expected chi-squared distribution with df = *n*-1, with *n* being the number of populations. Under a neutral evolution model, the loci are supposed to follow an expected chi-squared distribution if there is a single species.

We further investigated the global connectivity level between the MAG populations in the different Arctic Ocean regions. For this, the Arctic stations were divided based on their geographic location, as was previously done by (Royo-Llonch et al. 2021, p.). Five regions were identified: Pacific-Arctic, Kara-Laptev, Atlantic-Arctic, Arctic archipelago and Davis-Baffin (see Supplementary Figure S12 for the polar view of the Arctic *Tara* Oceans stations). To compare the level of genomic differentiation between these regions, median between regions pairwise-*F*ST values from all the *Chaetoceros* populations were extracted and their respective means compared.

### Estimation of variance partitioning

In a second step, the estimation of the relative effects of abiotic parameters and geographic distance on the genomic differentiation of MAGs was undertaken. For this, a linear mixed model from the R package ‘MM4LMM’ (Laporte and Mary-Huard 2021) was used as previously applied by Laso-Jadart et al. (2021) on marine plankton. As an input dataset for abiotic parameters, median values of different environmental parameters were extracted from the PANGAEA database (Ardyna et al. 2017; Guidi, Morin, et al. 2017; Guidi, Picheral, et al. 2017) and for each sampling site, namely oxygen, salinity, temperature, and nutrient concentrations of ammonium, iron, nitrate, nitrite, phosphate and silicate. Euclidean distances were then computed between the stations for all these parameters as well as for the station coordinates as a proxy for geographic distance. Finally, median pairwise-*F*ST values with the different abiotic parameters and distances were used as input for the LMM, allowing to estimate the relative proportion of the genomic variance explained by each parameter as well as an unexplained proportion. Mantel tests from the ‘vegan’ R package (Oksanen et al. 2020) were applied to verify the results.

### Identification of genes under selection

The ‘pcadapt’ R package v4.0.2 (Luu et al. 2017) was applied to detect selection among populations using the B-allele frequency (BAF) matrix, on ‘pool-seq’ mode with a minimal allele frequency of 0.05 within the populations, as was done in Laso-Jadart et al. (2021). Based on the PCA results, two samples from one station (194 SUR and DCM) of the MAG ARC_116 exhibited very distinct variation patterns (Supplementary Figure S24A), leading to a very high number of outliers (12,954) and were therefore removed from the analysis (Supplementary Figure 24B) to avoid false positive inflation. We computed q-values using the R package ‘qvalue’ with false discovery rate (FDR) correction (Storey et al. 2022). For the three MAGs ARC_116, ARC_217 and SOC_37, loci with a q-value < 0.15 were considered to be under selection. The functions of the genes harbouring loci under selection were investigated with the PFAM domains generated in part 3.5. Homologues of the genes of interest were subsequently searched by reverse genetics in *P. tricornutum* (Phatr3) by BLASTp. Manhattan plots of the contigs harbouring these loci were then built for each MAG, as well as bar plots representing their B-allele frequencies (BAF). To investigate the frequency of selected genes displaying loci under selection in other *Chaetoceros* species, searches of homologues among *Chaetoceros* spp. transcriptomes (derived from MMETSP; see the phylogeny section in Materials and Methods and Supplementary Figures S20 and S22) were performed by aligning target protein sequences on the 8 transcriptomes using tBLASTn. Output sequences were considered as homologue candidates when their scores and e-values were, respectively, at least equal to 200 and 1e-50. Alignments were conducted using the online version of MAFFT v7.463 (https://mafft.cbrc.jp/alignment/software/) (Katoh and Standley 2013). The sequences and alignments are available in the Supplementary Material online. Predictions of RNA secondary structures were conducted using RNAfold 2.4.18 (http://rna.tbi.univie.ac.at/) (Gruber et al. 2008) and Linearfold (beta) (http://linearfold.org/) (Huang et al. 2019) web servers.

## Acknowledgments

We would like to thank all colleagues from the *Tara* Oceans consortium as well as the *Tara* Ocean Foundation for their inspirational vision. This work was supported by the European Research Council (ERC) under the European Union’s Horizon 2020 research and innovation programme (Diatomic; grant agreement No. 835067). Additional funding is acknowledged from the French Government “Investissements d’Avenir” Programmes MEMO LIFE (Grant ANR-10-LABX-54), Université de Recherche Paris Sciences et Lettres (PSL) (Grant ANR-125311-IDEX-0001-02), France Génomique (ANR-10-INBS-09), and OCEANOMICS (Grant ANR-11-BTBR-0008). This article is contribution number *** of *Tara* Oceans.

## Author Contributions

C.N., A.M. and C.B. designed the study. E.P. retrieved and pre-processed the metagenomic data. C.N. performed the comparative genomics, phylogenetic analyses and genome-resolved metagenomics, interpreted the data, wrote the first manuscript draft and conceived the figures. C.N. performed the population genomic analyses with the support of A.M. C.N., A.M. and E.P. interpreted the data. All authors discussed the results and commented on the manuscript.

