## Supplementary_Figures for "Whole-genome scanning reveals selection mechanisms in epipelagic *Chaetoceros* diatom populations"

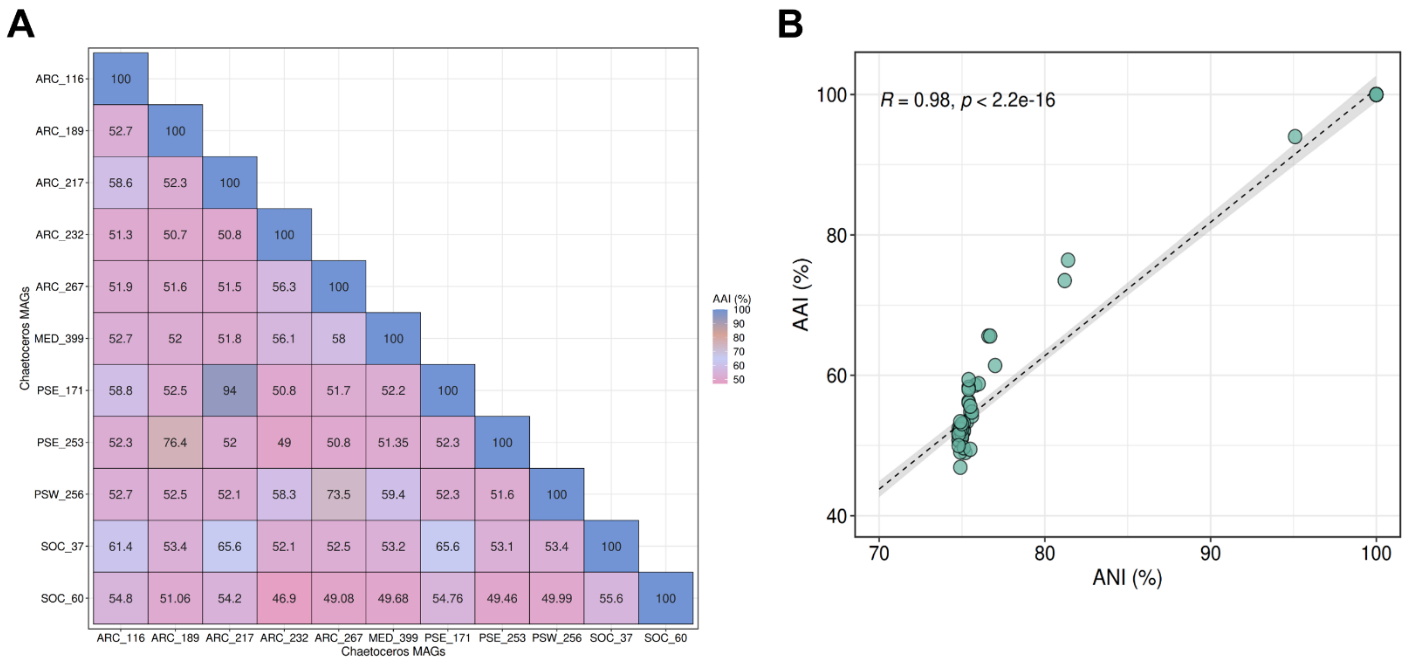


**Supplementary Figure S1.** (A) Average amino acid identity (AAI) of the MAGs. (B) Correlation analysis between ANI and AAI showing significant positive correlation (Pearson's correlation, the shaded area corresponds to 95% confidence interval).

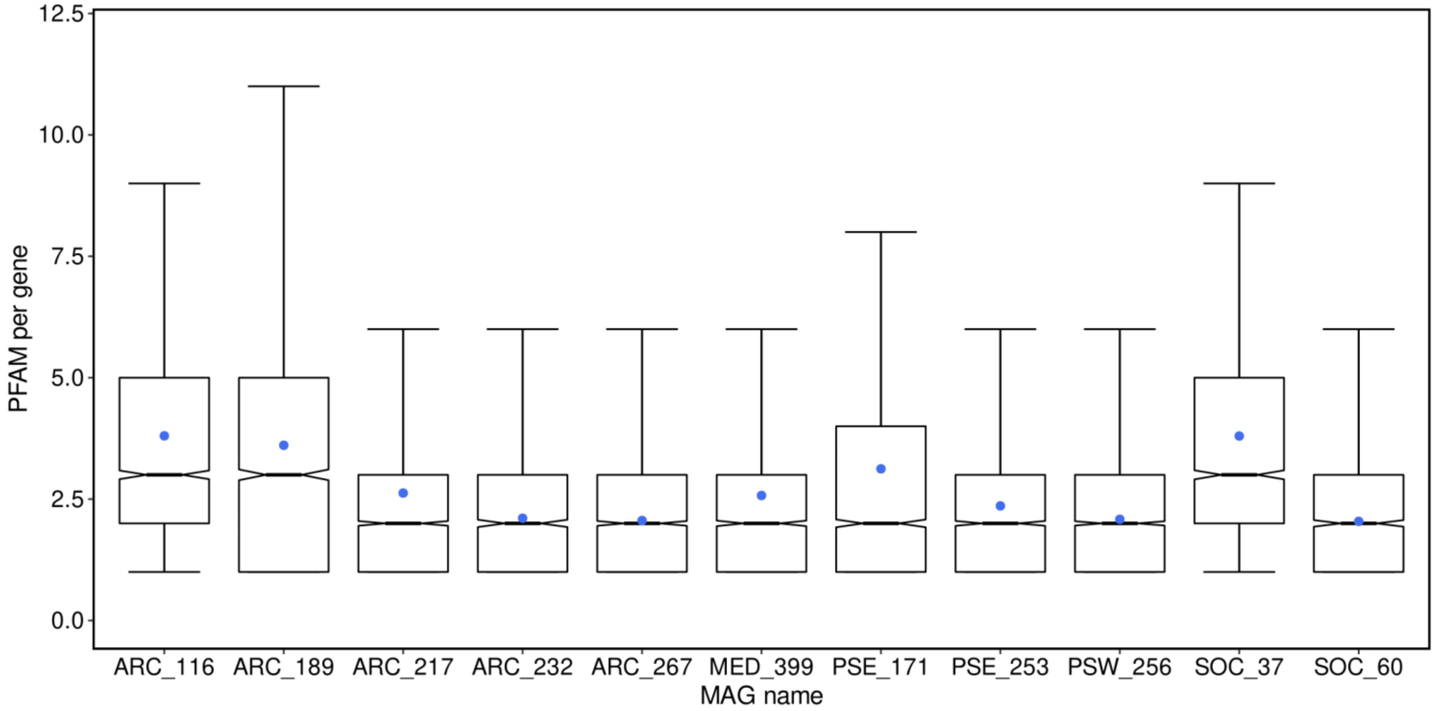


**Supplementary Figure S2.** Number of PFAM domains per gene. Boxplots plotting the distribution of the number of PFAM domains per gene with the blue dot representing the mean.

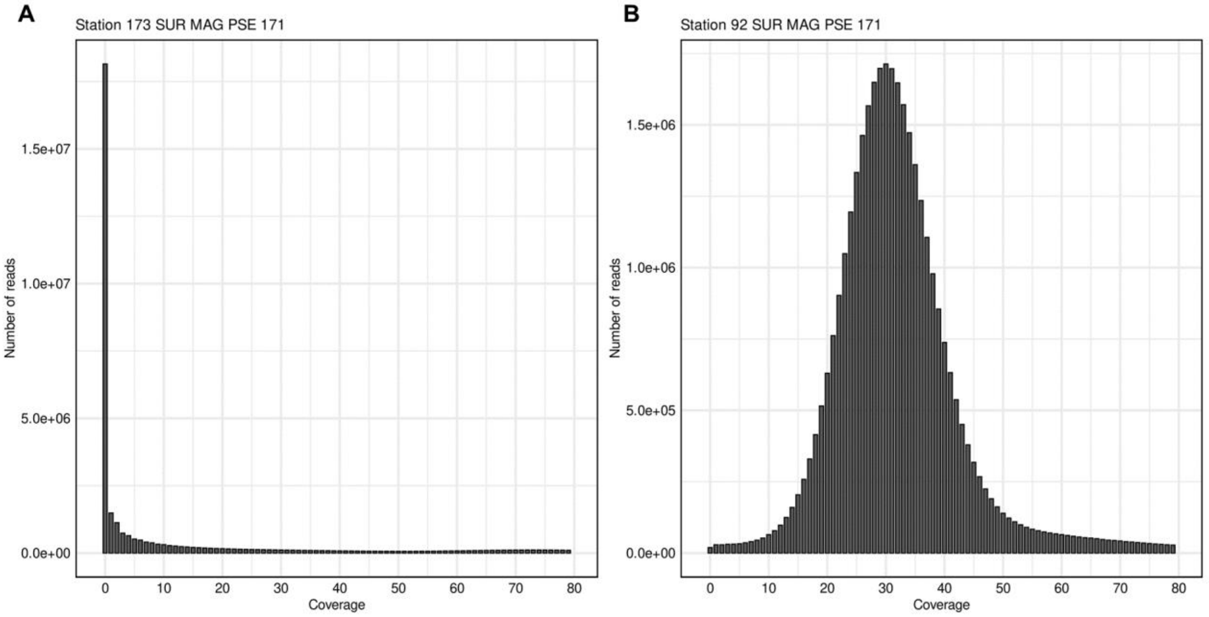


**Supplementary Figure S3.** Example of read coverage distributions. Read coverage distribution of MAG PSE_171 at the surface of stations (A) TARA_173 and (B) TARA_92. The distribution at station TARA_173 does not display enough coverage depth nor a clear unimodal pattern and will be discarded, whereas the pattern of TARA_92 is clearly unimodal and centered around 30X, hence the reads from this station will be kept for further analyses.

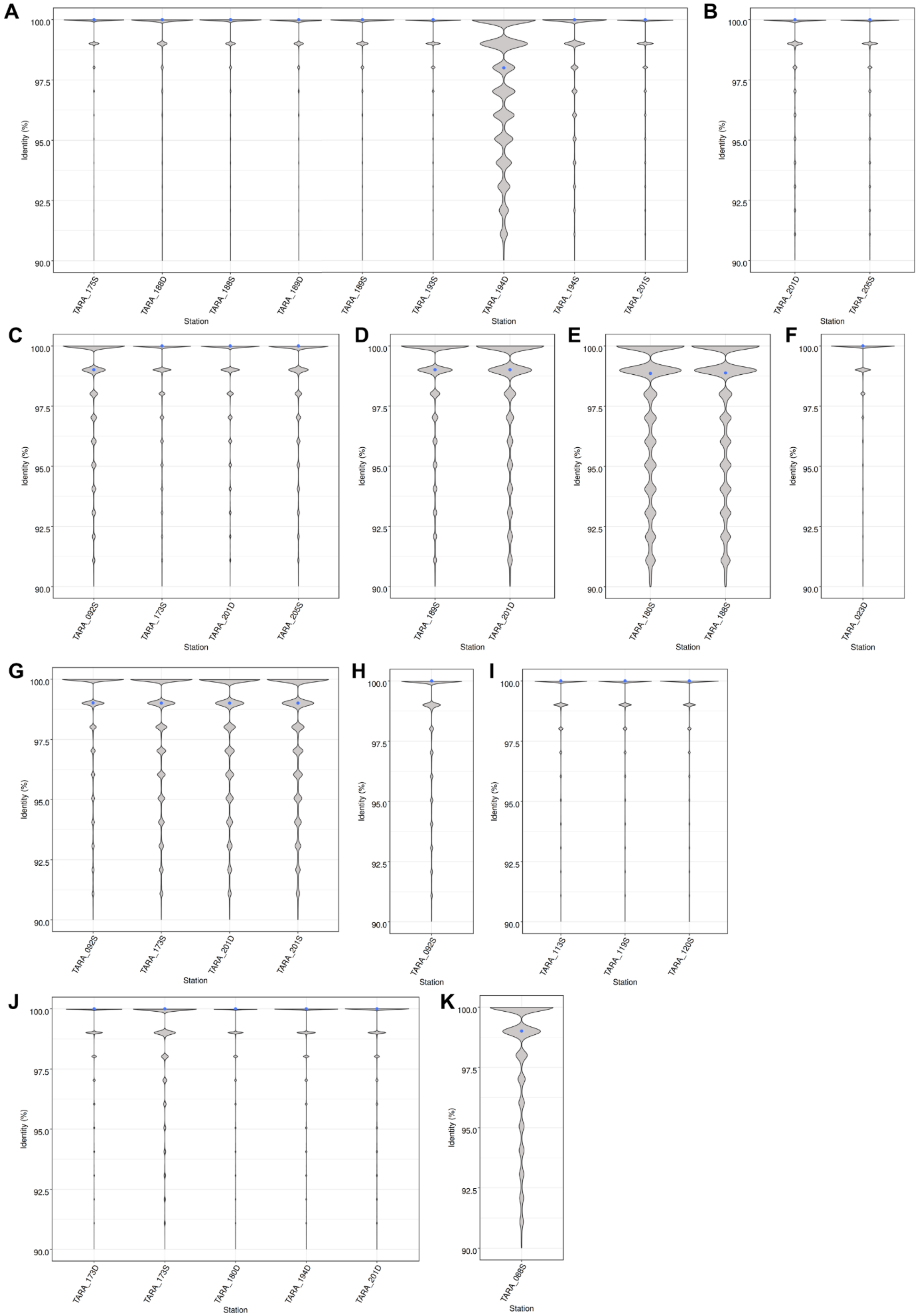


**Supplementary Figure S4.** Identity profiles of the metagenomic reads for the different *Chaetoceros* MAGs after filtration over 4x mean vertical coverage and 80% identity (plots shown starting at 90% identity for readability). (A) ARC_116, (B) ARC_189, (C) ARC_217, (D) ARC_232, (E) ARC_267, (F) MED_399, (G) PSE_171, (H) PSE_253, (I) PSW_256, (J) SOC_37 and (K) SOC_60. The blue dots represent mean identity.

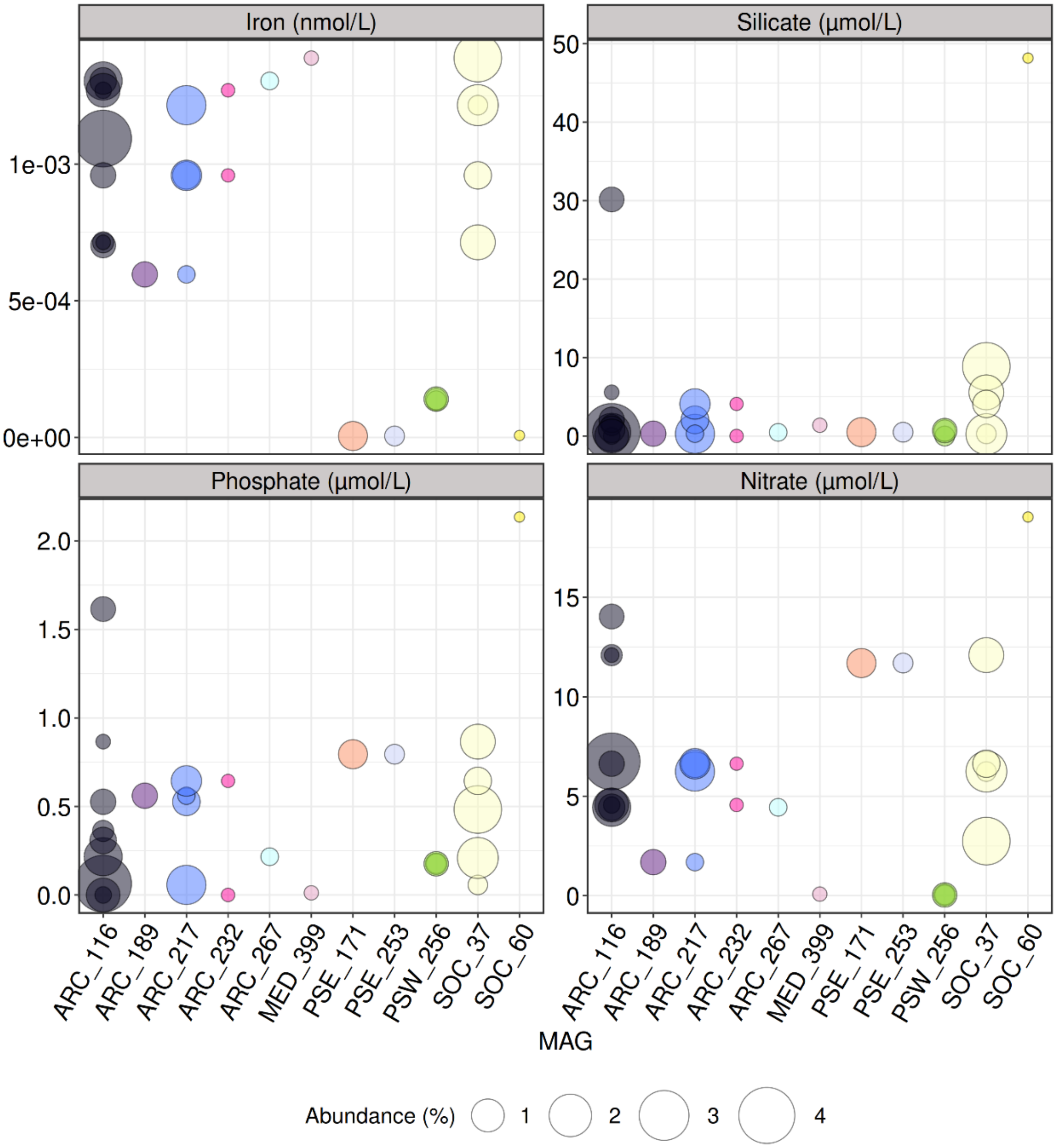


**Supplementary Figure S5.** Bubble plots of different nutrient concentrations at the respective sampling points of the MAGs. For a given MAG, most of the populations are distributed across a rather wide spectrum of iron, silicate, phosphate and nitrate concentrations.

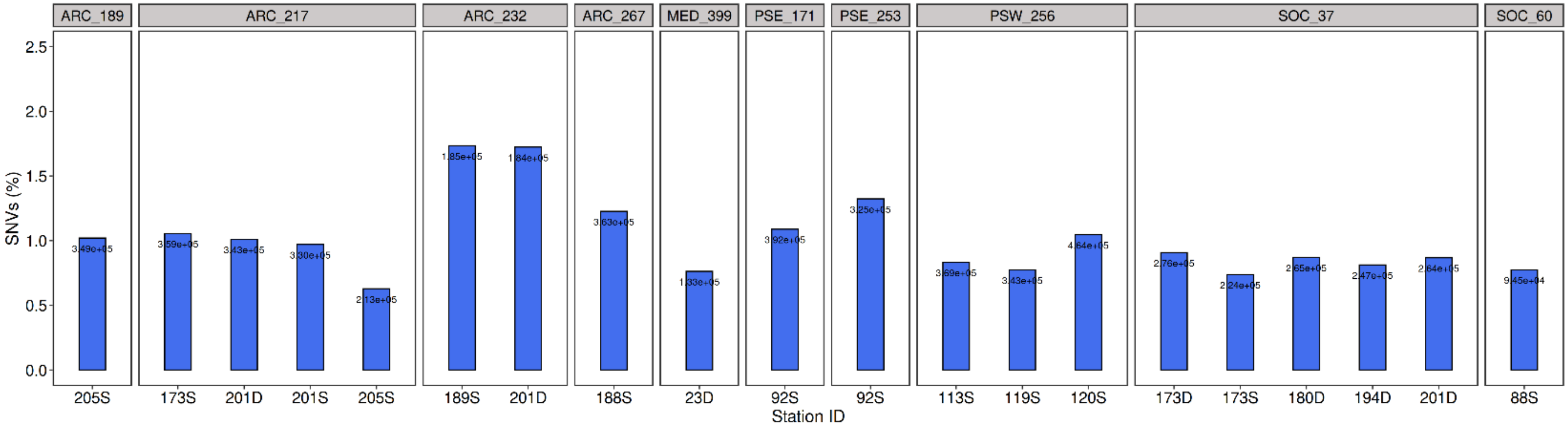


**Supplementary Figure S6.** Number of variants in the *Chaetoceros* populations. Single nucleotide variant (SNV) percentages are shown, together with the total number of variants.

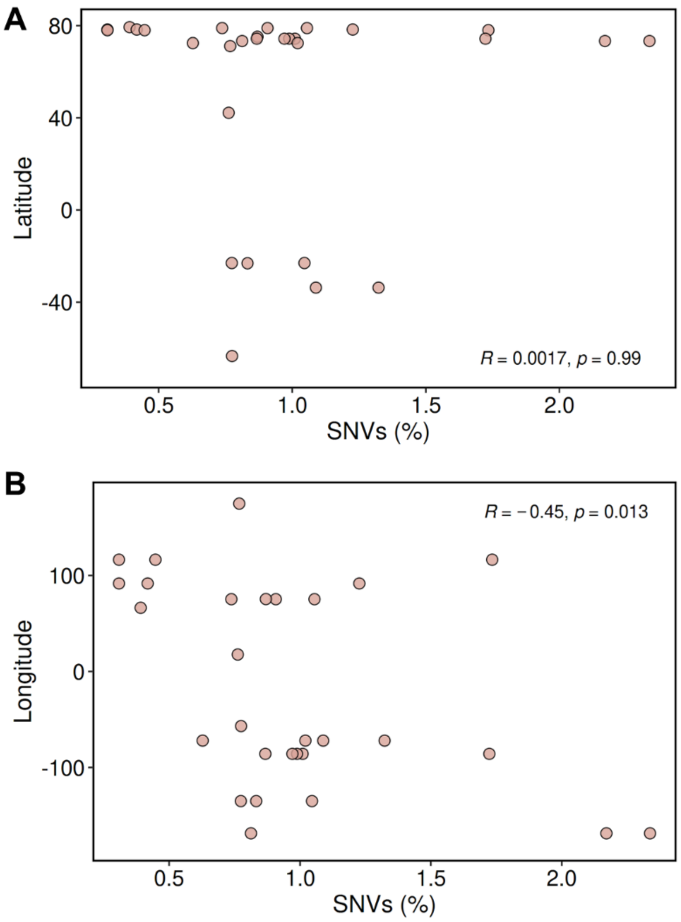


**Supplementary Figure S7.** Scatterplots showing the repartition of the SNVs in the populations regarding their (A) latitude and (B) longitude (Pearson's correlation coefficients and *p*-values are shown). No correlation was observed for the latitude while a negative effect of the longitude was observed.

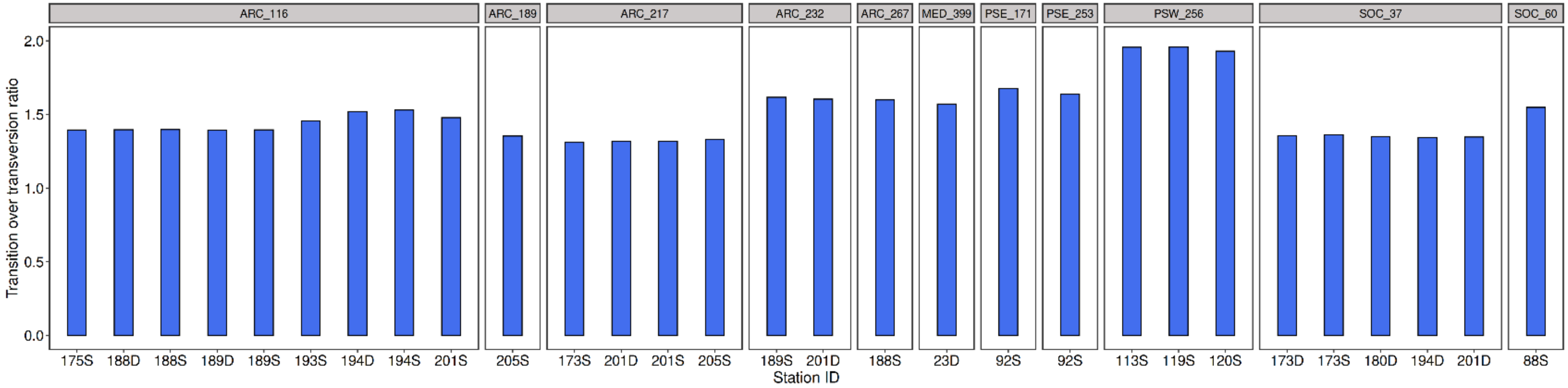


**Supplementary Figure S8.** Transition to transversion ratios of the *Chaetoceros* populations.

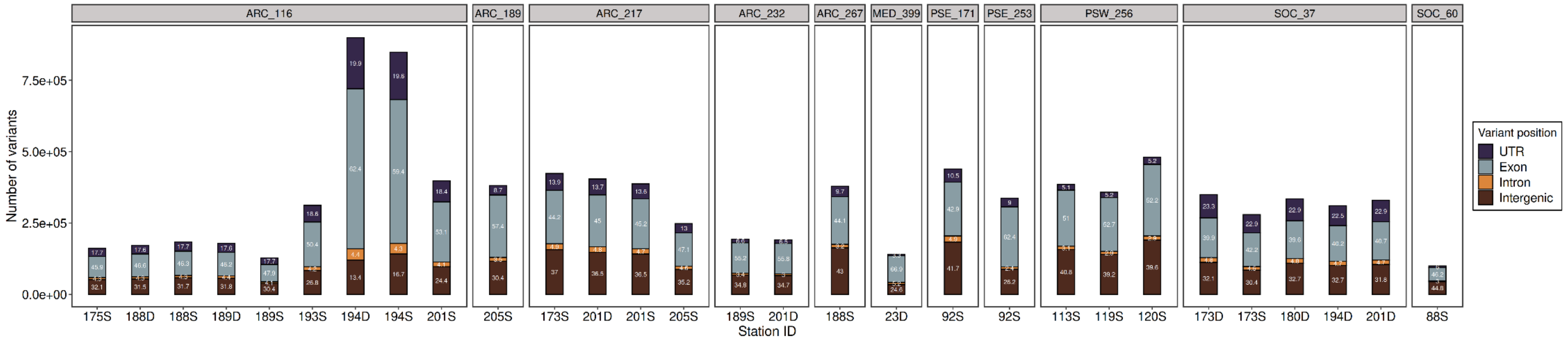


**Supplementary Figure S9.** Relative position of the SNVs in the *Chaetoceros* populations. The exact proportion of each category is annotated.

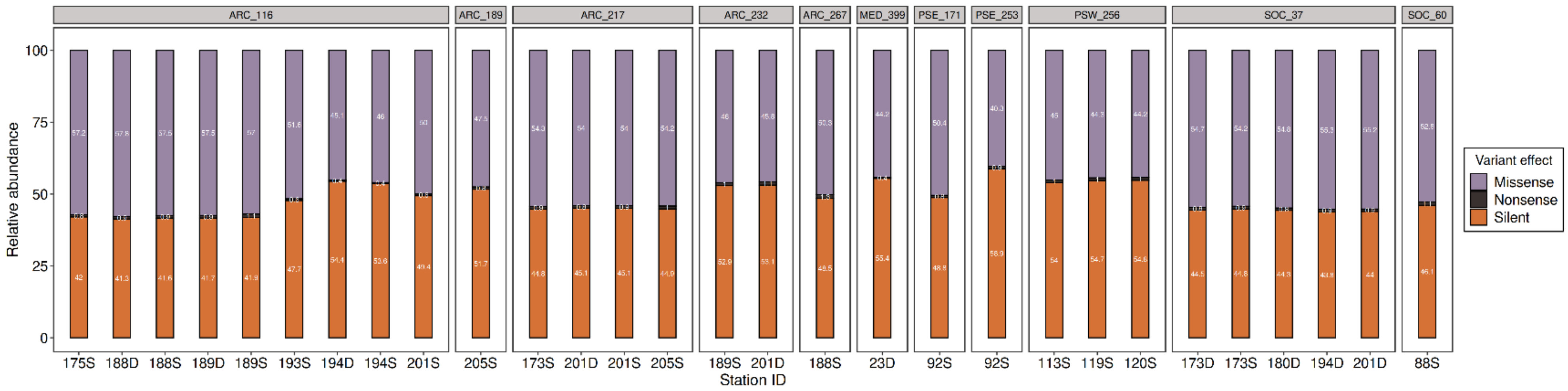


**Supplementary Figure S10.** Relative effects of the variants in the *Chaetoceros* populations.

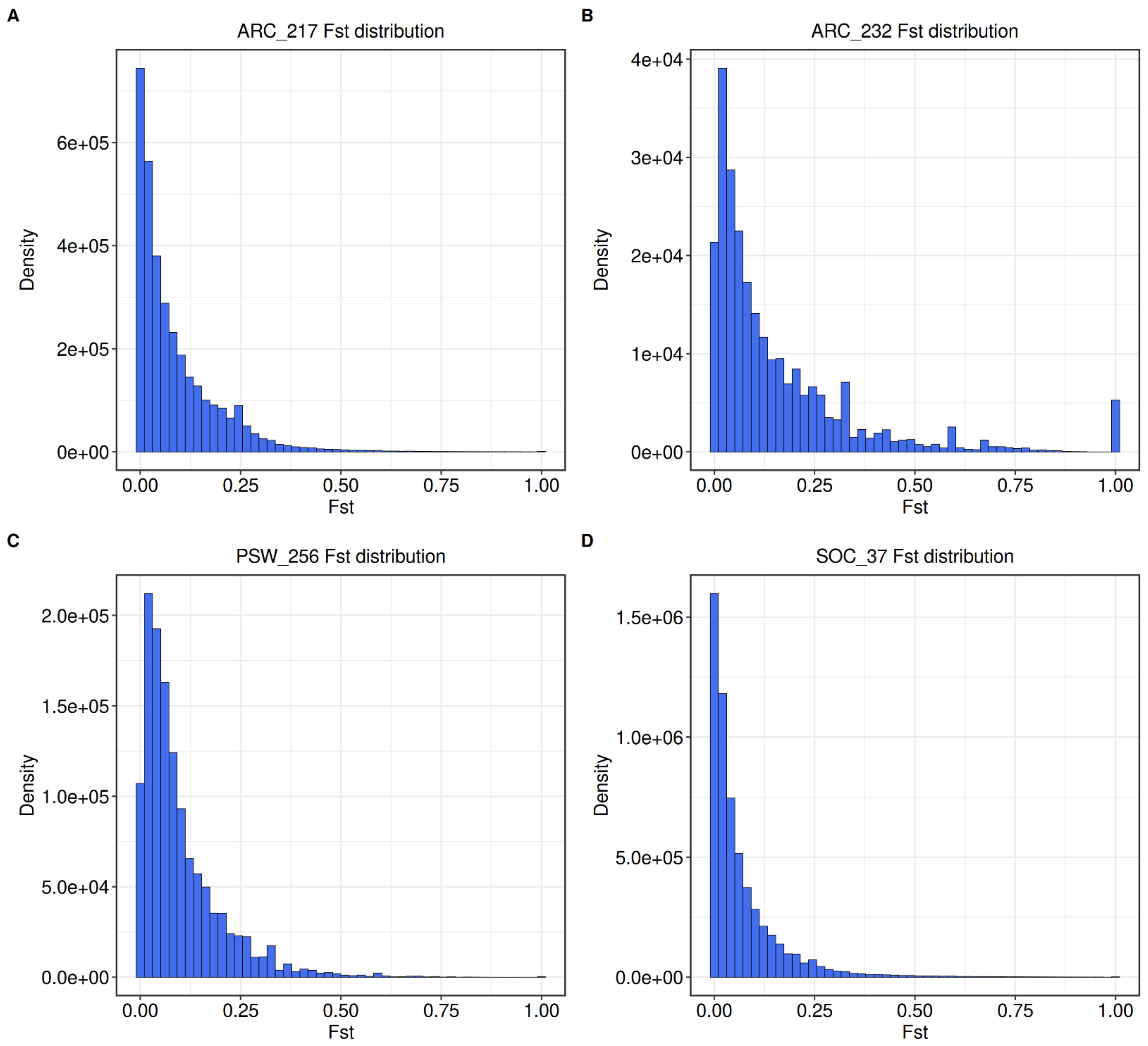


**Supplementary Figure S11.** Population-wide Fst profiles of the MAGs. The distributions are shown for the MAGs presenting at least two populations, that is (A) ARC_217, (B) ARC_232, (C) PSW_256 and (D) SOC_37. The Fst distributions appear globally unimodal, confirming that the reads of the respective MAGs were recruited from a single species.

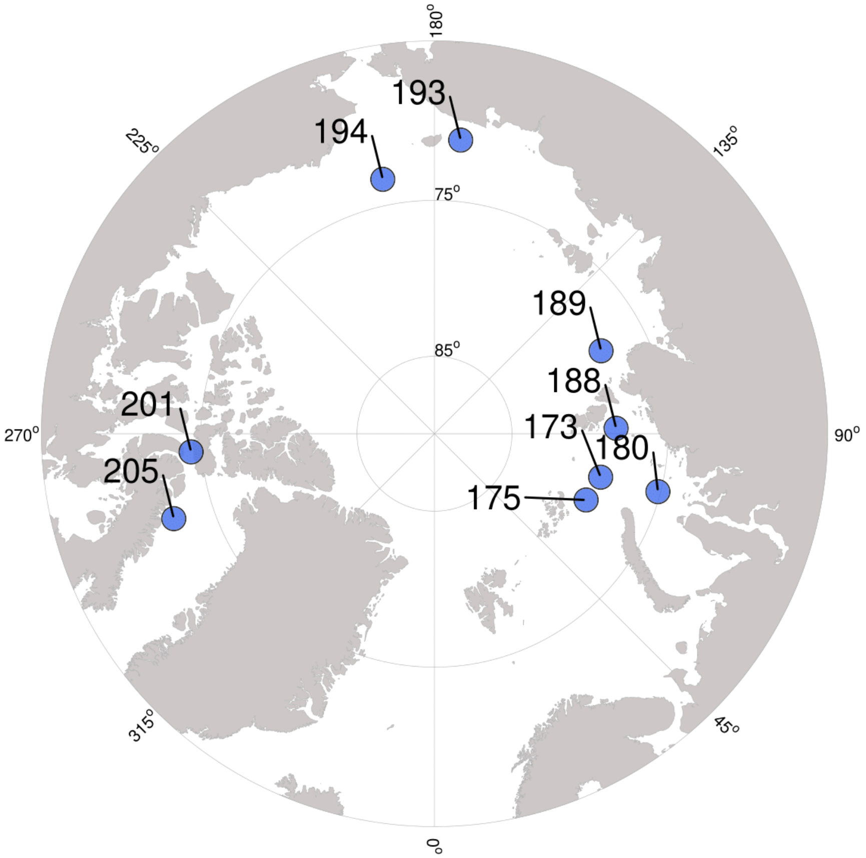


**Supplementary Figure S12.** Polar view of the *Tara* Oceans stations located in the Arctic Ocean where the *Chaetoceros* populations were identified. The *Tara* sampling stations belong to Arctic zones as follows: Atlantic-Arctic (175), Kara-Laptev (173, 180, 188 and 189), Pacific-Arctic (193 and 194), Arctic archipelago (201) and Davis-Baffin (205) (based on Royo-Llonch et al., 2021).

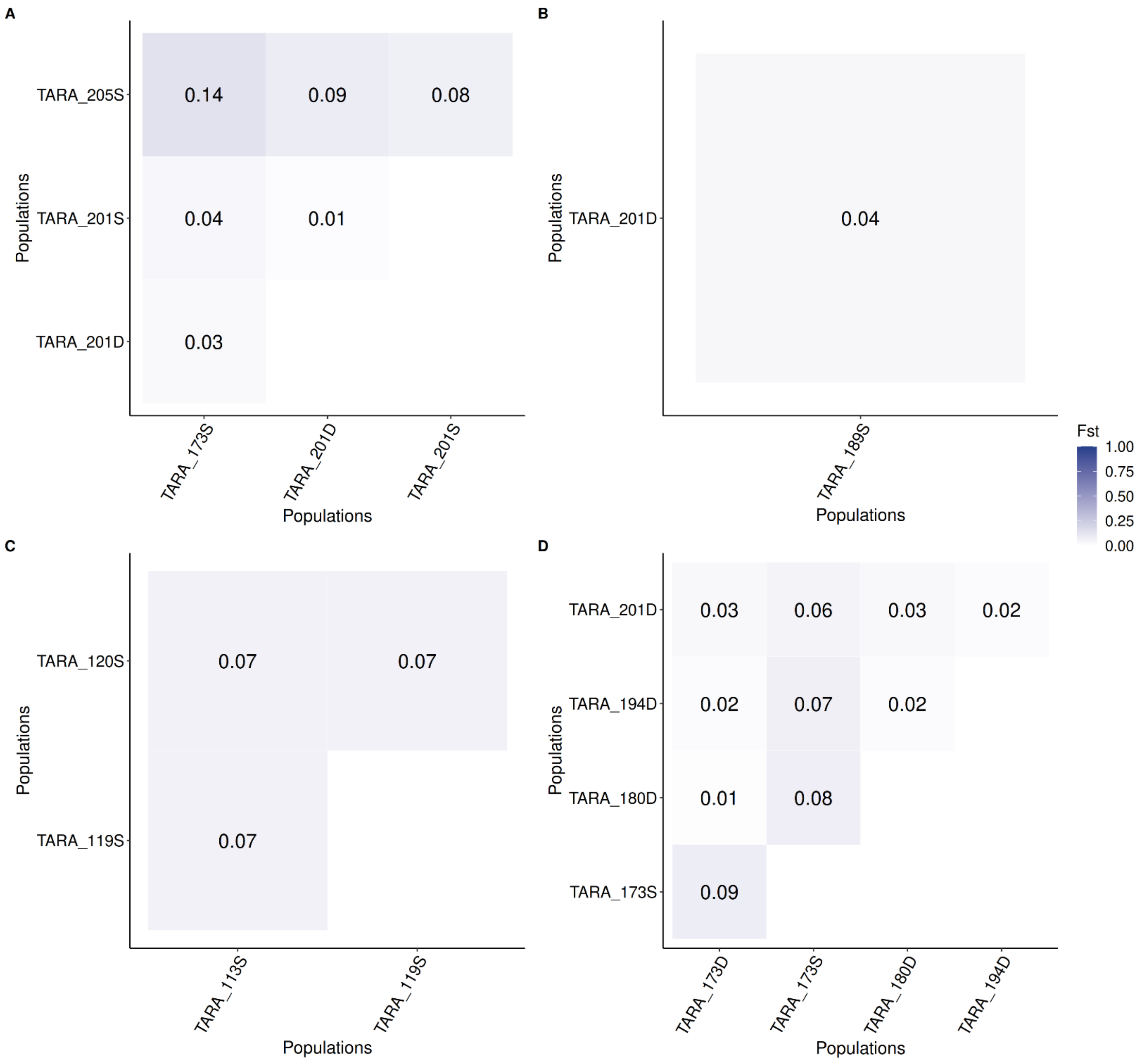


**Supplementary Figure S13.** Pairwise Fst matrices of the respective MAG populations. The matrices correspond to (A) ARC_217, (B) ARC_232, (C) PSW_256 and (D) SOC_37 populations.

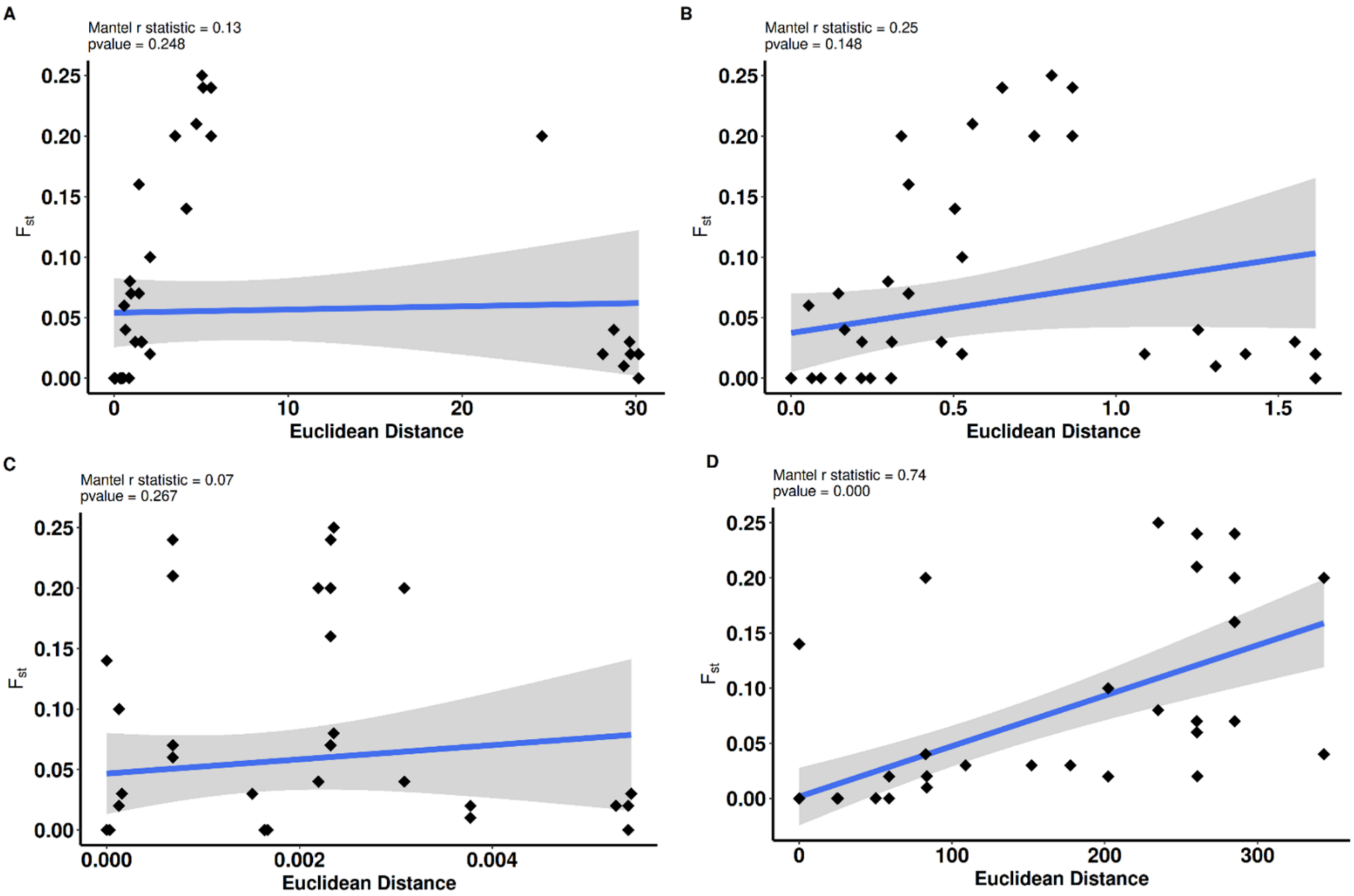


**Supplementary Figure S14.** Mantel tests of ARC_116 regarding the influence of (A) silicate, (B) phosphate, (C) nitrite and (D) geographic distance on the genetic differentiation.

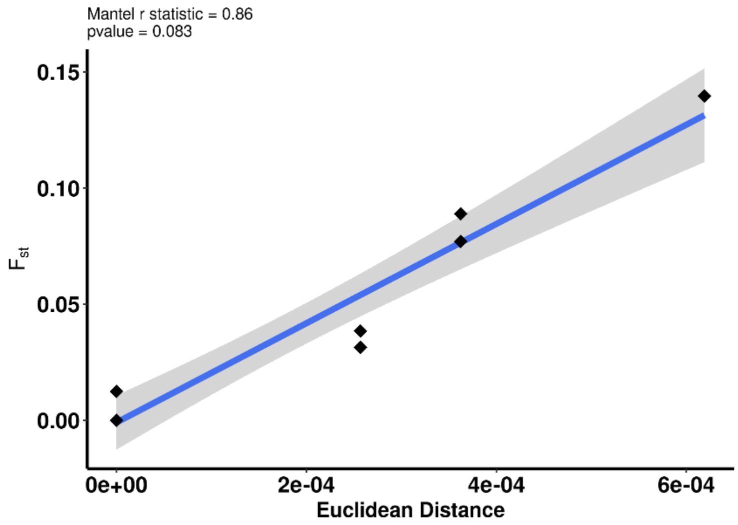


**Supplementary Figure S15.** Mantel test for ARC_217 regarding the impact of iron on the genetic differentiation.

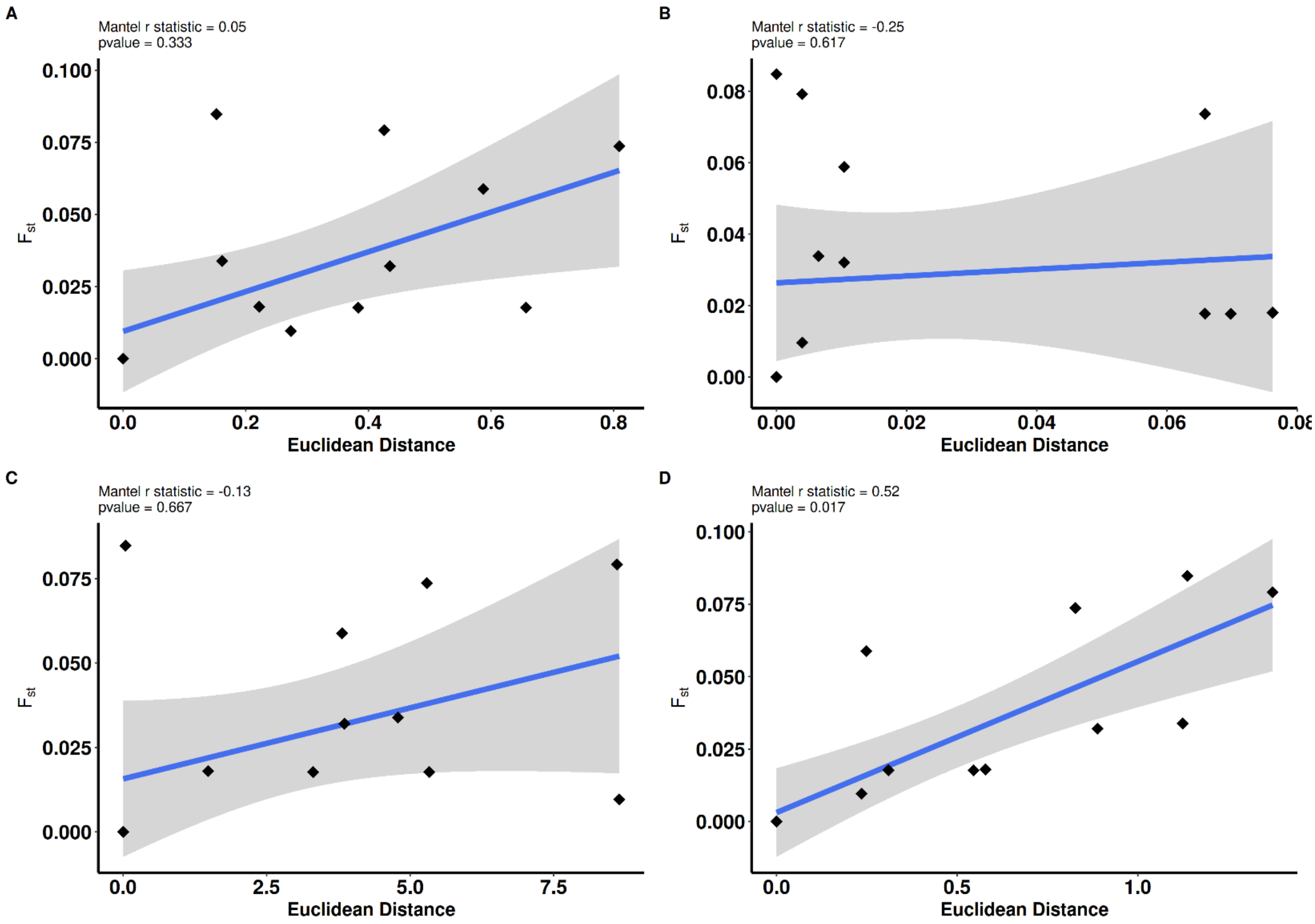


**Supplementary Figure S16.** Mantel tests of SOC_37 regarding the influence of (A) phosphate, (B) nitrite, (C) silicate and (D) temperature on the genetic differentiation.

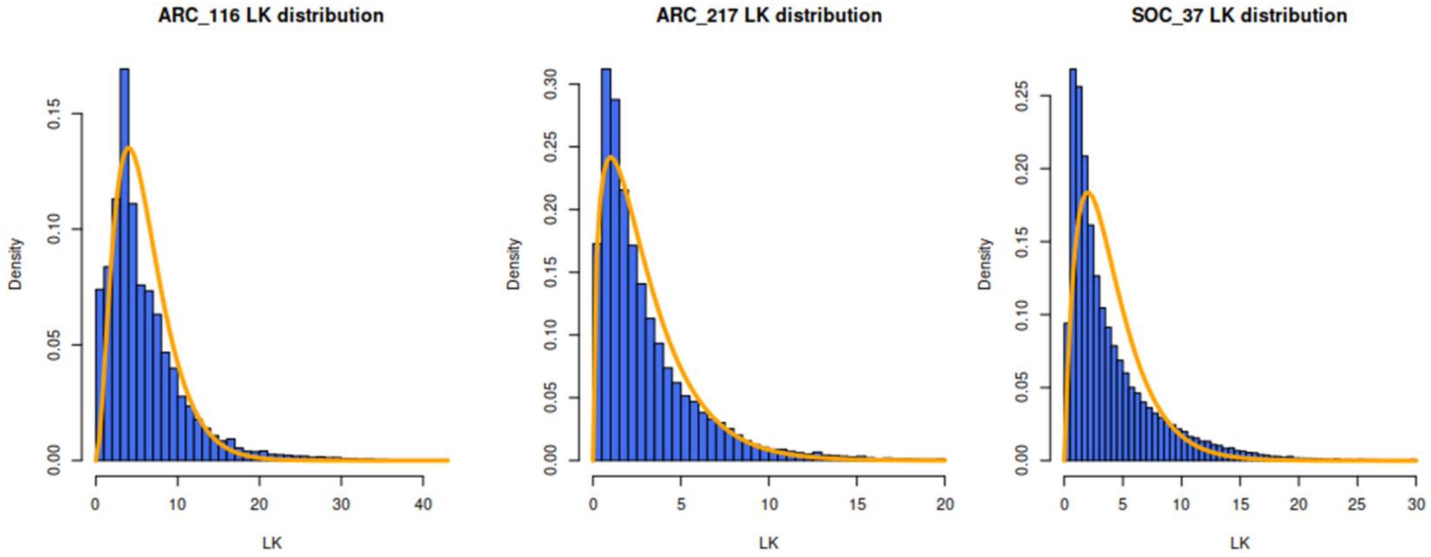


**Supplementary Figure S17.** Population-wide LK profiles of the MAG loci. The orange line indicates the χ2 (df=6; 3 and 4) theoretical distribution. The observed LK distributions followed the expected one.

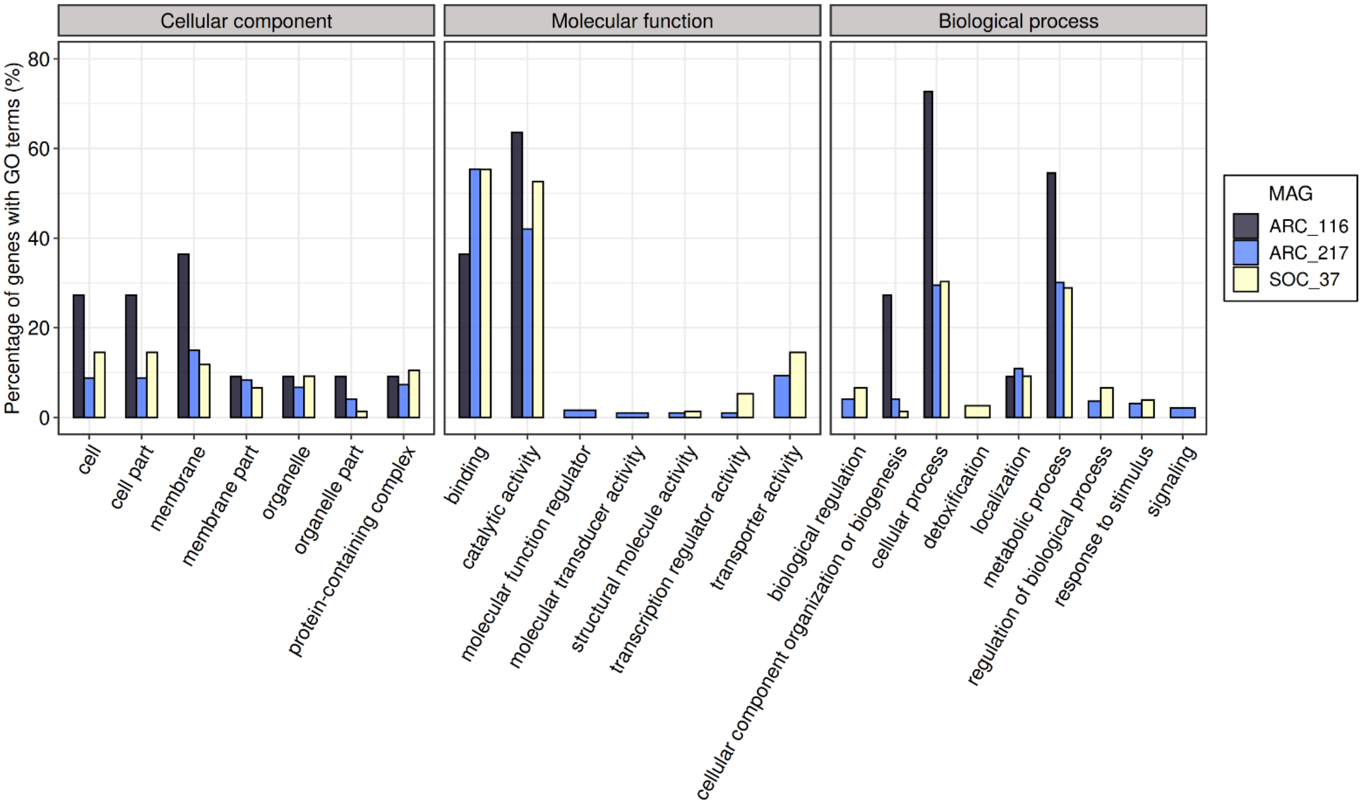


**Supplementary Figure S18.** GO terms associated with the loci under selection in *Chaetoceros* MAGs ARC_116, ARC_217 and SOC_37.

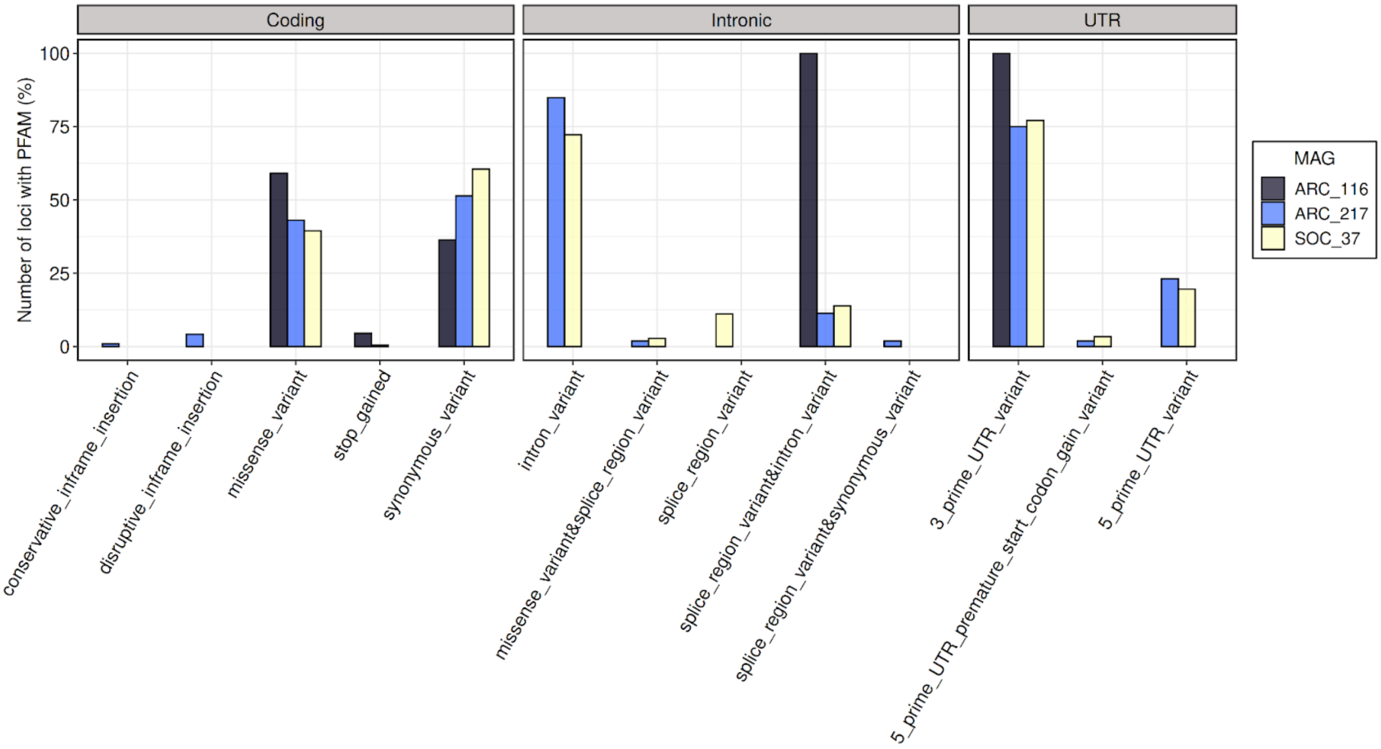


**Supplementary Figure S19.** Effect of the loci under selection within a PFAM domain, according to their position, for the MAGs ARC_116, ARC_217 and SOC_37.

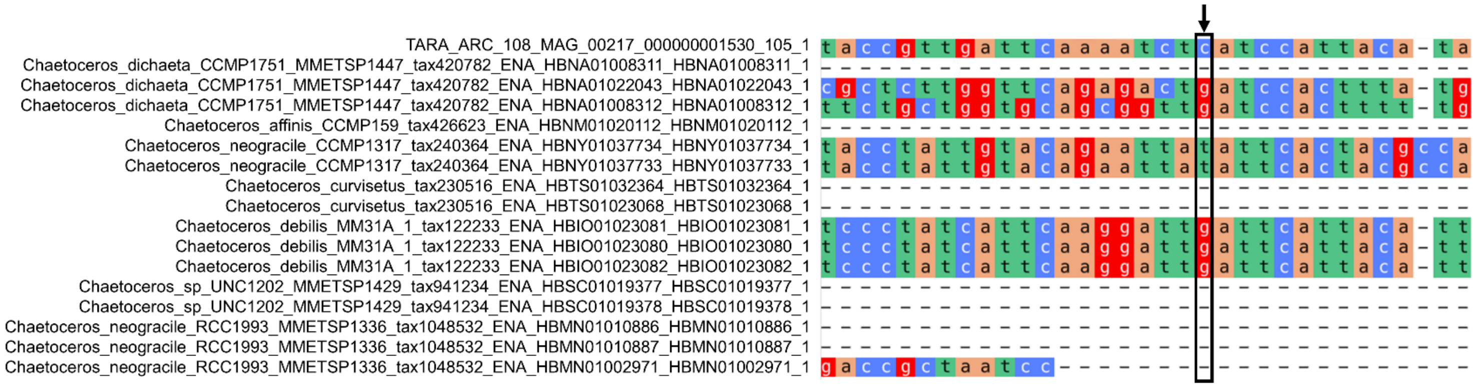

**Supplementary Figure S20.** Alignment of gene TARA_ARC_108_MAG_00217_000000001530.105.1 with homologue sequences from *Chaetoceros* spp. transcriptomes. Sequence alignment was realized with MAFFT v7 with the gene 1530.105.1 set as reference sequence. The position of the reference nucleotide undergoing selection in some ARC_217 populations is highlighted. The sequences and alignment are in Supplementary data files S1 and S2, Supplementary material online.

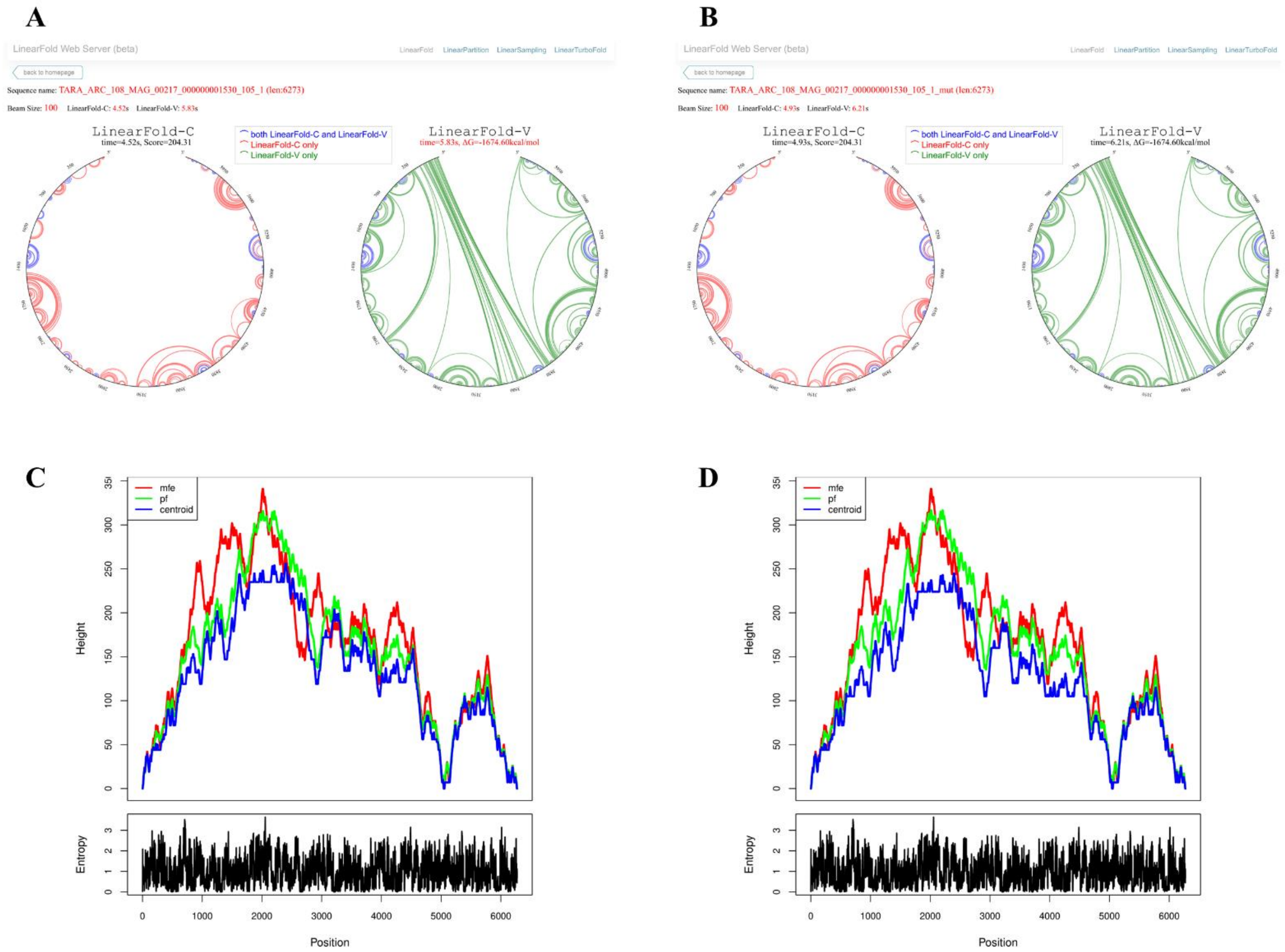


**Supplementary Figure S21.** RNA secondary structure predictions for gene TARA_ARC_108_MAG_00217_000000001530.105.1. Outputs from the LinearFold webserver for (A) the reference gene and (B) the gene with the p.Leu277Leu mutation showing exactly the same structure patterns. Mountain plots and positional entropy outputs from RNAfold representing the minimum free energy structure for the (C) reference gene and (D) gene with the SNV, displaying slightly different patterns.

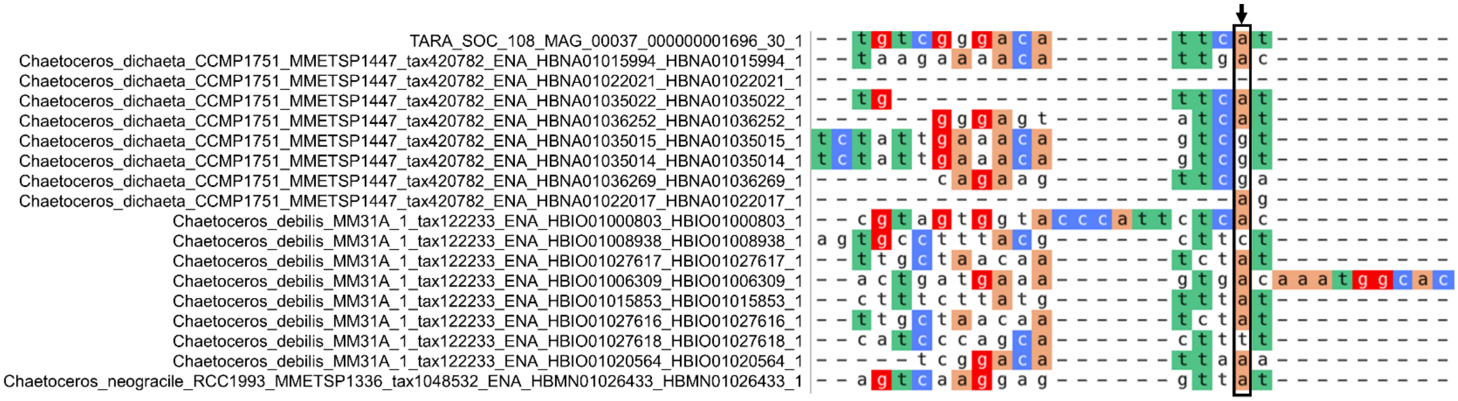

**Supplementary Figure S22**. Alignment of gene TARA_SOC_28_MAG_00037_000000001696.30.1 with homologue sequences from *Chaetoceros* spp. transcriptomes. Sequence alignment was realized with MAFFT v7 with the gene 1696.30.1 set as reference sequence. The position of the reference nucleotide undergoing selection in some SOC_37 populations is highlighted. The sequences and alignment are in Supplementary data files S3 and S4, Supplementary material online.

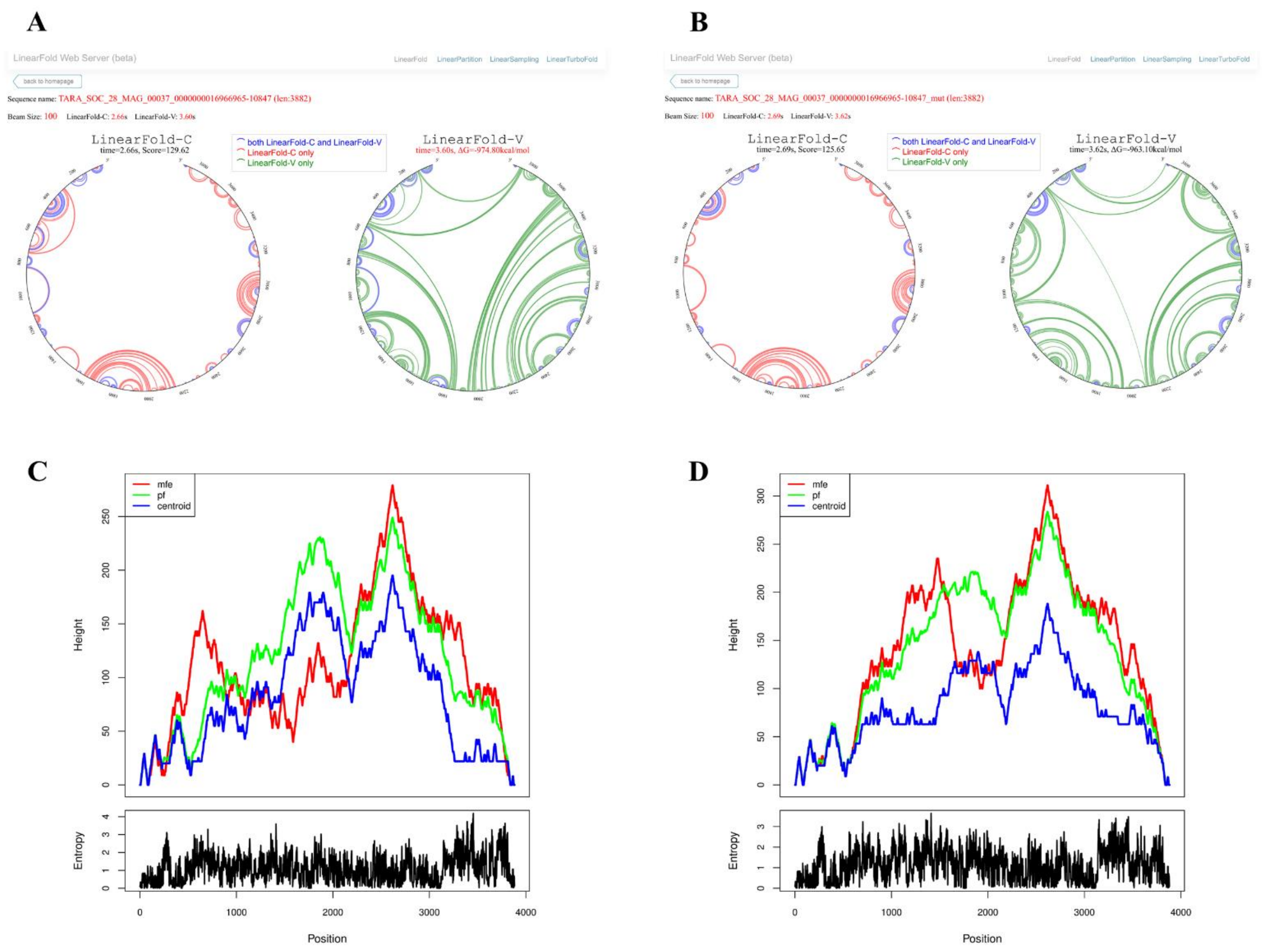


**Supplementary Figure S23.** RNA secondary structure predictions for gene TARA_SOC_28_MAG_00037_000000001696.30.1. Outputs from the LinearFold webserver for (A) the reference gene and (B) the gene with the p.Met1063Leu mutation showing significantly different structure patterns. Mountain plots and positional entropy outputs from RNAfold representing the minimum free energy structure for the (C) reference gene and (D) gene with the SNV, displaying again different patterns.

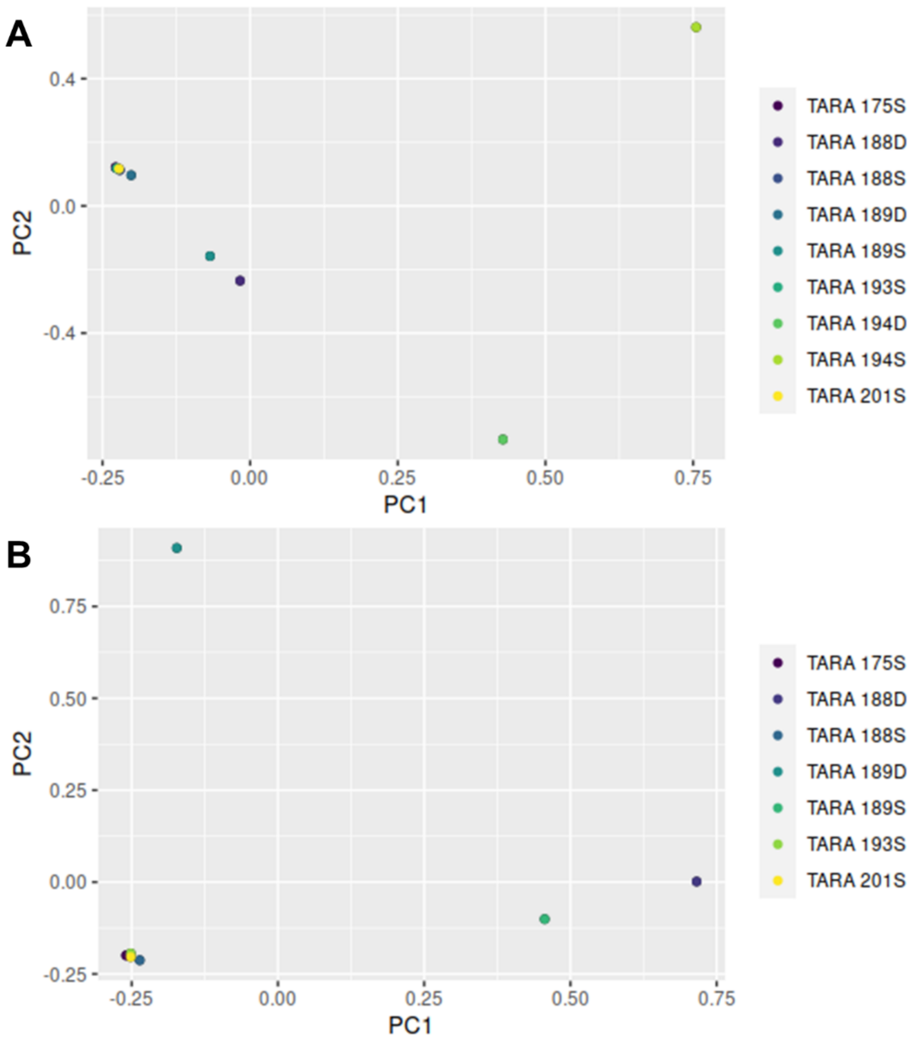


**Supplementary Figure S24.** Principal component analysis of ARC_116 allelic frequencies in the different populations. (A) The PCA of all the stations shows that populations from TARA_194 at the surface and DCM are pulled away from the others. (B) The PCA after removing the outlier points from station TARA_194 shows more details on the allelic frequency patterns, with less heterogeneous dispersion. Stations TARA_188D, TARA_189S and TARA_189D are clearly apart from the others.
